# NREM consolidation and increased spindle counts improve age-related memory impairments and hippocampal representations

**DOI:** 10.1101/2020.01.20.912915

**Authors:** Robin K. Yuan, Matthew R. Lopez, Manuel-Miguel Ramos-Alavarez, Marc E. Normandin, Arthur S. Thomas, David S. Uygun, Vanessa R. Cerda, Amandine E. Grenier, Matthew T. Wood, Celia M. Gagliardi, Herminio Guajardo, Isabel A. Muzzio

## Abstract

Age-related changes in sleep patterns have been linked to cognitive decline. Specifically, increasing age is associated with increasing fragmentation of sleep and wake cycles. However, it remains unknown if improvements in sleep architecture can ameliorate cellular and cognitive deficits. We evaluated how changes in sleep architecture following sleep restriction affected hippocampal representations and memory in young and old mice. After training in a hippocampus-dependent object/place recognition task, control animals were allowed to sleep *ad libitum*, while experimental animals underwent 5 hours of sleep restriction (SR). Interestingly, old SR mice exhibited successful object/place learning comparable to young control mice, whereas young SR and old control mice did not. Successful learning correlated with the presence of two hippocampal cell types: 1) “Context” cells, which remained stable throughout training and testing, and 2) “Object” cells, which shifted their preferred firing location when objects were introduced to the context and moved during testing. As expected, EEG analysis revealed more fragmented sleep and fewer spindles in old controls than young controls during the post-training sleep period. However, following the acute SR session, old animals exhibited increased consolidation of NREM and increased spindle count, while young mice only displayed changes in REM bout length. These results indicate that consolidation of NREM sleep and increases in spindle count serve to ameliorate age-related memory deficits and allow hippocampal representations to adapt to changing environments.

**eTORC Blurb:** Age-related cognitive decline is associated with poor sleep quality. This study shows that acute sleep restriction serves to improve memory, hippocampal representations, and sleep quality in old mice, having the opposite effect in young animals. These findings indicate that improving sleep quality may mitigate age-related cognitive decline.

**Highlights:** Acute sleep restriction improves memory in old mice, but adversely affects young ones

Acute sleep restriction makes hippocampal representations more flexible in old mice

Acute sleep restriction improves sleep quality and increases spindle count in old mice

Acute sleep restriction decreases hippocampal flexibility in young mice

## Introduction

A large amount of evidence suggests that sleep plays a key role in memory consolidation (Abel et al., 2013; Rasch and Born, 2013; Stickgold and Walker, 2013; Tononi and Cirelli, 2014). Studies have shown performance gains following post-training sleep (Gais and Born, 2004; Gais et al., 2006; Smith, 2001), as well as learning impairments when sleep restriction (SR) is conducted after training (Graves et al., 2003; Prince et al., 2014; Smith and Rose, 1996). However, the effects of SR are complex and vary across the lifespan. For example, partial SR has minimal effects on adolescent cognitive performance (de Bruin et al., 2017) and sleep loss can differentially impact adults (Krause et al., 2017), with some studies showing within- and across-individual differences in cognitive susceptibility (Saletin et al., 2016; Wilson et al., 2019). Interestingly, SR therapy, which is characterized by limiting sleep periods, has very positive effects on the sleep quality of old subjects (Wennberg et al., 2013), yet the underlying physiological and cellular changes associated with this improvement remain unclear.

Sleep involves interspersed periods of non-rapid eye movement (NREM) sleep, a state characterized by high amplitude, low frequency (0.2-4 Hz), synchronous electroencephalographic (EEG) activity, and rapid eye movement (REM) sleep, which is characterized by low amplitude fast desynchronized EEG waves. It has been proposed that NREM is particularly important for memory formation because during this stage information is transferred to cortex for long-term storage (for a review see, (Antony et al., 2019), whereas REM has been associated with both consolidation of novel information and forgetting of previously encoded information (Poe, 2017). During NREM, there are rapid bursts of activity (10-14 Hz) of short duration, known as spindles. Spindles are thought to facilitate memory reactivation, which is essential for proper consolidation (Rasch and Born, 2013). Critically, changes in NREM and spindle characteristics predict early memory impairments in older subjects (Taillard et al., 2019). However, it is unknown if reversing these spindle changes can have positive effects on cognition.

The hippocampus plays a critical role in the formation of episodic memories – recollections of events happening in specific contexts at particular times (Smith and Mizumori, 2006; Squire and Zola, 1998). The activity of hippocampal place cells, which fire in specific locations when animals navigate (O’Keefe and Dostrovsky, 1971), provides a cognitive map in which episodic memories are embedded (Mizumori, 2006; Smith and Mizumori, 2006). Evidence supporting this idea comes from the observation that networks of cells active during wake are reactivated in similar sequences/patterns during NREM sleep at compressed time scales (Drieu et al., 2018; Hwaun and Colgin, 2019; Lee and Wilson, 2002), which has been shown to be important for goal-directed memories (de Lavilleon et al., 2015).

Both sleep and cognition have been shown to undergo age-related changes across the lifespan (Huang et al., 2002). Older humans exhibit more fragmented sleep and less slow wave sleep in comparison to younger adults (Espiritu, 2008; Hasan et al., 2012; Ohayon et al., 2004). Additionally, older subjects exhibit impairments in hippocampus-dependent cognitive tasks (Lester et al., 2017; Lister and Barnes, 2009; Miller and O’Callaghan, 2005). These observations have suggested that age-related cognitive decline may be linked to changes in sleep patterns (Altena et al., 2010); however, the relationship between sleep quality, cognitive performance, and hippocampal activity during wake periods remains unclear. Here, we investigated if an acute period of SR could modify subsequent sleep architecture, hippocampal place cell firing, and object-place recognition (OPR) memory in young and old mice. Our results indicate that SR has differential effects in young and old mice. In young mice, an acute SR session does not significantly affect sleep architecture but it does decrease the flexibility of hippocampal representations and memory. Conversely, in old animals, SR leads to NREM consolidation and increases in spindle count, which allows the hippocampus to adapt to changing environments and reverses age-related memory deficits.

## Results

### Sleep restriction impairs object-place recognition memory in young adult mice but enhances performance in old mice

Animals were trained in the OPR task as illustrated in **Figure 1A**. The analysis of object/place preference during the test showed an interaction between age and sleep condition [Fw(1)=0.55, p<0.00001]. Simple effects indicated that the young controls displayed greater preference for the moved object in comparison to young SR [young control (YC) vs. young SR (YSR); Tw(16.07)=3.39, p<0.02]. Conversely, old SR mice displayed greater preference for the moved object compared to old controls [old control (OC) vs. old SR (OSR): Tw(15.10)= 2.11, p<0.05, **Figure 1B**].These results suggested that SR had a surprising beneficial effect on OPR memory in old animals, bringing their performance to the level of young controls.

**Figure 1.**
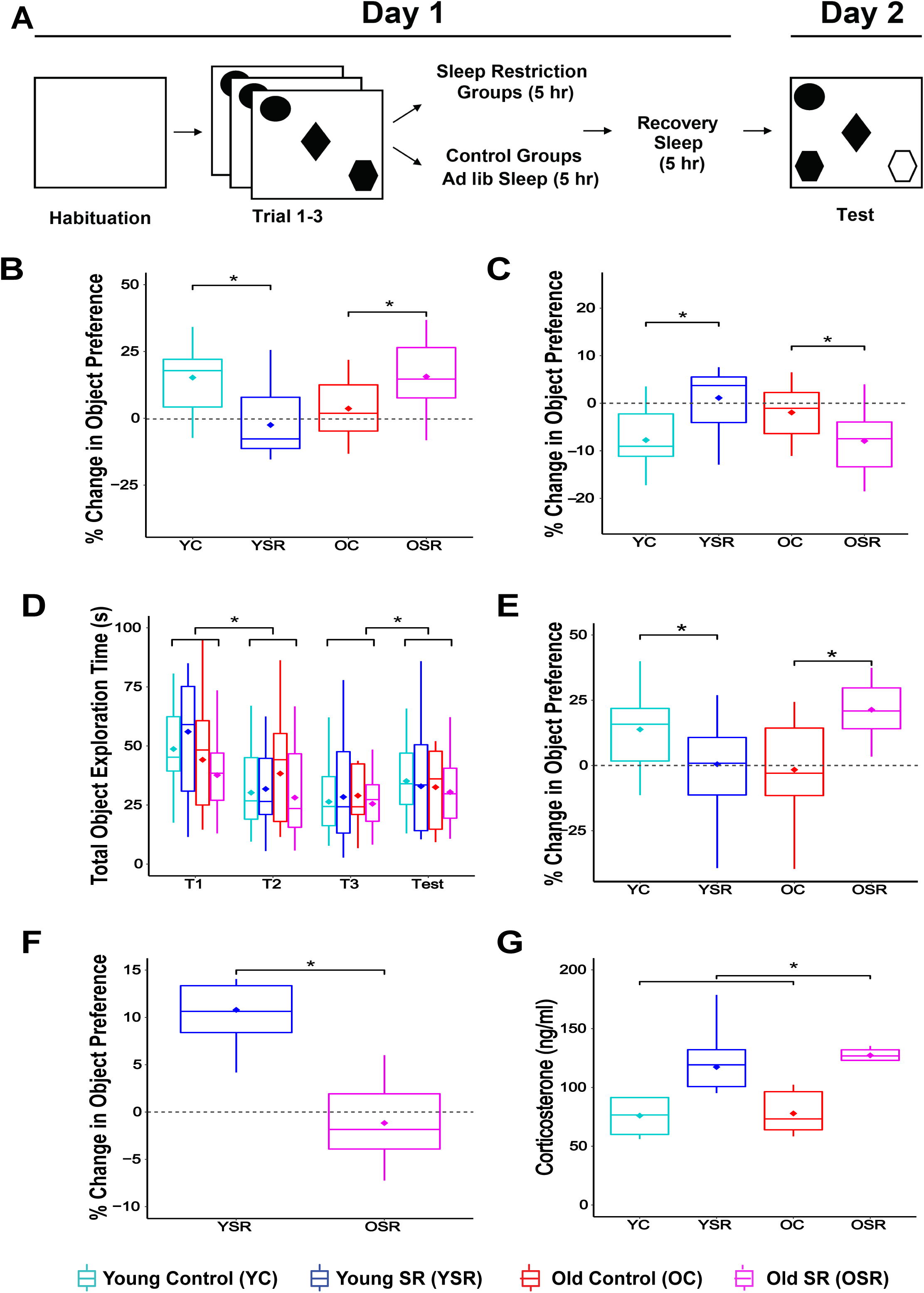
Object place recognition (OPR) behavioral performance. A) Schematic of behavioral design. B) Percent change in preference for displaced object [number of animals (n): young control (YC):n=16, young SR (YSR): n=15, old control (OC): n=13, old SR (OSR): n=15]. C) Percent change in preference for the unmoved objects. D) Total time of object exploration during training and testing in seconds (s). E) Percent change in preference for displaced object excluding trial 1. F) Percent change in preference for displaced object in young (n=7) and old (n=7) mice receiving SR 5 hr after training (delayed SR, YSR: n=6, OSR: n=7). G) Levels of plasma corticosterone in control (young: n=5; old: n=5) and SR (young: n=5; old: n=6) mice. In all Figures data are graphed using Box Plots according to robust statistics based on the median. The Box- Plot describes the data through their quartiles. The inner line corresponds to the median, the lower box represents the first quartile, the upper box the third quartile, and extending lines, known as whiskers, represent the minimum and maximum, showing the extent of variability outside the lower and upper quartiles. The diamond represents the trimmed mean (e.g., adjusted average without outliers) used for analysis. T1-T3: training trials. Asterisks (*) represent significance using alpha=0.05. Statistical details in Supplemental Tables.

Experiments with rats revealed that animals display more exploration of familiar objects when the animals perceive the context as different (Cohen et al., 2013; Dix and Aggleton, 1999; Mumby et al., 2002). To determine if failures in context recognition modulated preference for familiar object location, we calculated the percent change in preference for the unmoved objects. We found an interaction between age and sleep condition [Fw(1)=0.55, p<0.0001, **Figure 1C**]. Single effects corroborated that young controls and old SR mice displayed less preference for the unmoved objects [YC vs. YSR: Tw(16.07)=11.51, p<0.01, OC vs. OSR: Tw(15.10)=4.47, p<0.05], but there were no significant differences between the young SR and old controls [YSR vs. OC: Tw(15.58)=1.30, p=0.26]. These data indicate that the poor OPR performance exhibited by young SR and old controls is not related to a shift in context-dependent preference for objects placed in familiar locations.

We evaluated the total time exploring the objects across trials to confirm that there were no group differences in locomotor activity. We found no significant effects of age, sleep condition, or interactions on total exploration [age: Fw(1)=0.12, p=0.73; sleep: Fw(1)=0.07, p=0.79, interaction (age vs. sleep): Fw(1)=0.31, p=0.38; interaction (age vs. sleep vs. trial): Fw(3)=0.09, p=0.81]. However, there was a trial effect [Fw(3)=4.08, p<0.0001]. Simple effects revealed that all groups displayed more total object exploration during the first object trial as well as during the test [T1-T2: Zw=3.82, p<0.0001; T3-Test: Zw=1.75, p<0.04, **Figure 1D**], which likely resulted from the initial novelty of the objects and the change in configuration, respectively. Since we observed this difference, we then calculated the percent change in object preference excluding the first object trial to ensure that our results were not biased by a novelty effect. The results were almost identical to those including all trials [interaction age vs. sleep: Fw(1):0.55, p<0.0001, Young: control vs. SR: p<0.05; old: control vs. SR: p<0.003; **Figure 1E**]. These findings demonstrated that the differences in object preference observed across the groups were not due to variability in object exploration.

The observation that old SR animals displayed better OPR performance suggested that post-learning SR followed by recovery sleep had a beneficial effect on memory. To determine if this effect was time-dependent, we delayed the SR by 5 hours in a separate group of young and old animals. These mice were trained and tested exactly as described above but were allowed to sleep *ad libitum* for 5 hours after training, prior to the SR procedure. This manipulation did not impair young mice, which showed a preference for the moved object during the test in comparison to training. Additionally, it did not rescue the age-dependent memory impairments in old animals, which displayed no object preference during the test [Tw(6.88)=4.85, p<0.002, **Figure 1F]**. These results indicated that there is a time window during which SR can produce memory deficits or ameliorate age-related cognitive decline in young and old animals, respectively.

### Levels of corticosterone are similar in young and old sleep restricted mice

Total or partial sleep restriction (SR) has been shown to induce mild stress in rodents (Coenen and van Luijtelaar, 1985; Tobler et al., 1983). Additionally, sleep alterations correlate with a rise in cortisol levels in older human subjects (Castello-Domenech et al., 2016; Morgan et al., 2017). We collected blood from young and old mice following 5 hours of SR (experimental groups) or 5 hours of *ad libitum* sleep (control groups) to determine if our method had a differential effect on corticosterone levels in our groups. In agreement with previous findings in C57Bl/6 lines (Kolbe et al., 2015; Oh et al., 2018), we observed slightly more variability in the young animals than old ones. Critically, although SR produced an increase in plasma corticosterone compared to normal sleep conditions [F(1)=16.85, p<0.02]; there were no differences between the age groups [F(1)=0.29, p=0.61] or interactions between age and sleep condition [F(1)=0.04, p=0.87, **Figure 1G**]. These data indicate that corticosterone levels cannot account for the differences in performance between young and old SR mice.

### Rate remapping increases during testing in young control and old sleep deprived animals

A subset of animals was implanted with tetrodes in area CA1 to determine the effects of sleep restriction on hippocampal neuronal activity during OPR performance (electrode positions are shown in Supplemental Figure 1A). First, we corroborated that behavioral performance of tetrode-implanted animals paralleled the results obtained in animals with EMG/EEG headmounts (Table **1**). Then, we examined place cell activity in 110 cells recorded in 10 young mice (60 cells in 6 controls and 50 cells in 4 SR mice) and 77 cells recorded in 10 old mice (36 cells in 5 controls and 41 cells in 5 SR mice). There were no significant effects of age or group by trial interactions in mean, peak, or out of field firing rate [p>0.05, Supplemental Figure 1B-D]. Additionally, there were no differences or interactions in number of fields, field size, or spatial information content (p>0.05, Supplemental Figure 1E-G). These data indicate that place cell parameters did not differ between age groups or sleep conditions throughout training and testing.

**Table 1.** Performance of tetrode and EEG/EMG implanted animals in the OPR task. Behavior of tetrode and EEG/EMG implanted animals

|  | Young Control | Young SR | Old Control | Old SR |
| --- | --- | --- | --- | --- |
| All animals | 14.04 ± 2.82 | 0.05 ± 3.56 | 1.69 ± 2.76 | 16.79 ± 3.43 |
| Tetrode animals | 15.45 ± 5.83 | 2.09 ± 6.76 | -4.39 ± 10.19 | 18.23 ± 7.36 |
| EMG/EEG animals | 13.27 ± 3.38 | -0.83 ± 4.35 | 5.50 ± 3.46 | 16.07 ± 3.89 |

Hippocampal cells have previously been shown to code environmental changes through increases or decreases in firing rate, a phenomenon known as rate remapping (Leutgeb et al., 2005). Therefore, we hypothesized that the analysis of average firing activity could have masked potential rate remapping differences. To assess this possibility, we calculated rate remapping as the absolute difference in peak firing rate for each cell across trials. This analysis revealed a main effect of age group [Fw(1)=10.20, p<0.00001], an effect of trial [Fw(3)=2.12, p<0.005] and an interaction between age, trial, and sleep condition [Fw(3)=1.30, p<0.05]. Analysis of simple effects indicated that rate remapping was different between the groups exclusively during the test trial. Young controls and old SR animals, the two groups that exhibited successful learning, displayed more rate remapping than young SR during the test session [YC vs. YSR: Tw(46.29)=2.30, p<0.03, YSR vs. OSR: Tw(29.79)=3.93, p<0.005] and increased rate remapping during the test in comparison to the last training trial (T3 vs. Test: YC: Zw=1.69, p<0.02; OSR: Zw=2.80, p<0.02, **Figure 2A**). These results indicate that changes in rate remapping are important for updating object memory representations in the OPR task.

**Figure 2.**
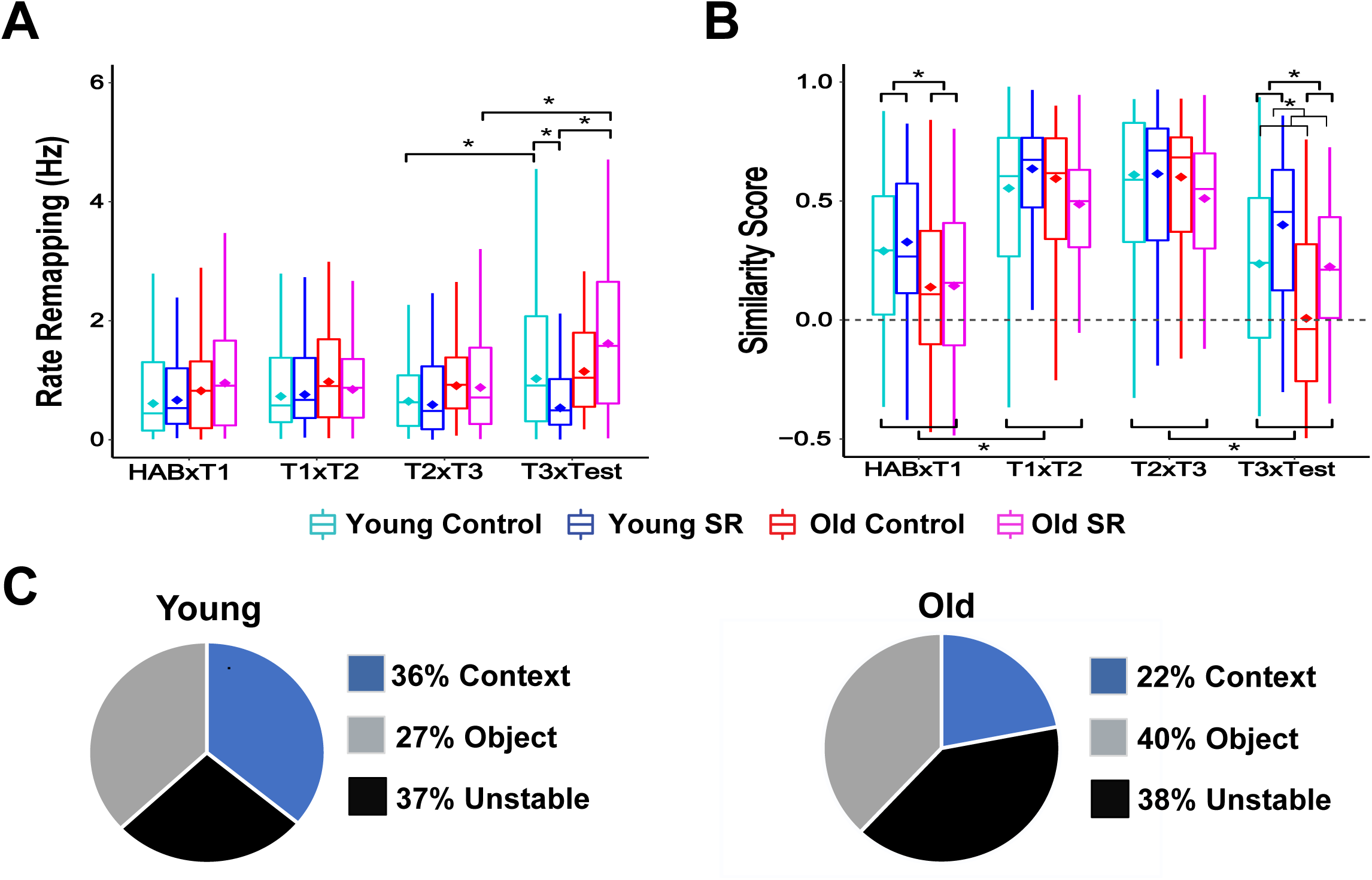
Rate and global remapping across sessions and percentage of different cell types during OPR performance. A) Young controls and old SR mice displayed higher rate remapping during the test trial in comparison to the last training trial (T3). Additionally, these groups also displayed more rate remapping than young SR mice during the test. B) Average global remapping (map similarity) for all groups across trials. Old animals displayed significant more instability than young mice between the habituation and trial 1 and during testing. C) Percentage of context, object, and unstable cells recorded in young (right panel) and old left panel) mice. Young controls: 6 mice, 60 cells, range of cells per animal: 6-16, young SR: 4 mice, 50 cells, range of cells per animal: 8-18, old control: 5 mice, 36 cells, range of cells per animal: 5-13, old SR: 5 mice, 41 cells: range of cells per animal: 4-13. Hab: habituation, T1-T3: training trials. Asterisks (*) represent significance using alpha=0.05. Statistical details in Supplemental Tables.

### The overall stability of hippocampal representations decreases during the moved-object test in all groups

Hippocampal cells respond to spatial cues, including objects (Cohen et al., 2013). Therefore, we expected a shift the cells’ preferred firing locations (i.e. global remapping) when the objects were first introduced (Hab vs. T1), but short-term stability across the training trials when the objects and environment remained unchanged (T1 to T3). We also anticipated global remapping during the test trial (Test), reflecting the change in the object configuration. Analysis of similarity between place cell maps revealed a significant effect of age [Fw(1)=8.32, p<0.008], trial [Fw (3) =29.35, p<0.000001], and interactions of age and trial [Fw(3)=1.45,p<0.04] and sleep condition and trial [Fw(3)=1.63,p<0.02]. Analysis of simple effects showed that old mice displayed more global remapping than young animals when the objects were first introduced [Tw(98.60)=2.95, p<0.004], which likely reflected the unstable nature of spatial representations in this group (Barnes et al., 1997). However, no differences in stability were observed during the object training trials [T1 vs. T2: Tw(101.10)=1.08, p=0.28, T2 vs. T3: Tw(95.97)=1.12, p=0.26]. As expected, all groups displayed lower place field stability during the test trial in comparison to the high short-term stability observed during training (p<0.05), with old animals displaying more instability than the young ones [T3 vs. test: Tw(91.84)=3.40, p<0.001, **Figure 2B]**. These results indicate that all groups exhibit some global remapping during the moved object test trial.

### Distinct cell types are differentially affected by sleep restriction and age

We have previously observed that distinct CA1 subpopulations respond differently during learning (Wang et al., 2015). To determine if the same happened during the OPR task, cells were classified according to their remapping patterns during training. “Context” cells displayed high stability throughout habituation and training; “object” cells remapped when objects were first introduced but remained stable during all the object training sessions; and “unstable” cells remapped across all training sessions. In young animals, each of these categories made up roughly 1/3 of the cells recorded. However, old animals had a lower percentage of stable context cells than young mice (**Figure 2C**).

After cells were classified, we examined similarity scores (e.g., the degree of global remapping) within each group across consecutive trials (**Figure 3A**). We hypothesized that if animals remembered the environment, context cells should remain stable during the moved- object test. Context cells showed an effect of age [Fw(1)=7.87, p<0.007], a main effect of trial [Fw(3)=3.04, p<0.0001], and an interaction between age, sleep condition, and trial [Fw(3)=0.82, p<0.05]. Analysis of simple effects showed that context cells in old control animals were more unstable than in old SR or young controls [OC vs. OSR: Tw(7.30)=2.82, p<0.03; YC vs. OC: Tw(7.38)= 3.31, p<0.02]. Interestingly, there was no difference between young controls and young SR mice in the stability of context cells [Tw(20.06)=1.03, p=0.32, **Figure 3B**], suggesting that the memory impairment observed in the latter may not be due to a failure in recalling the environment.

**Figure 3.**
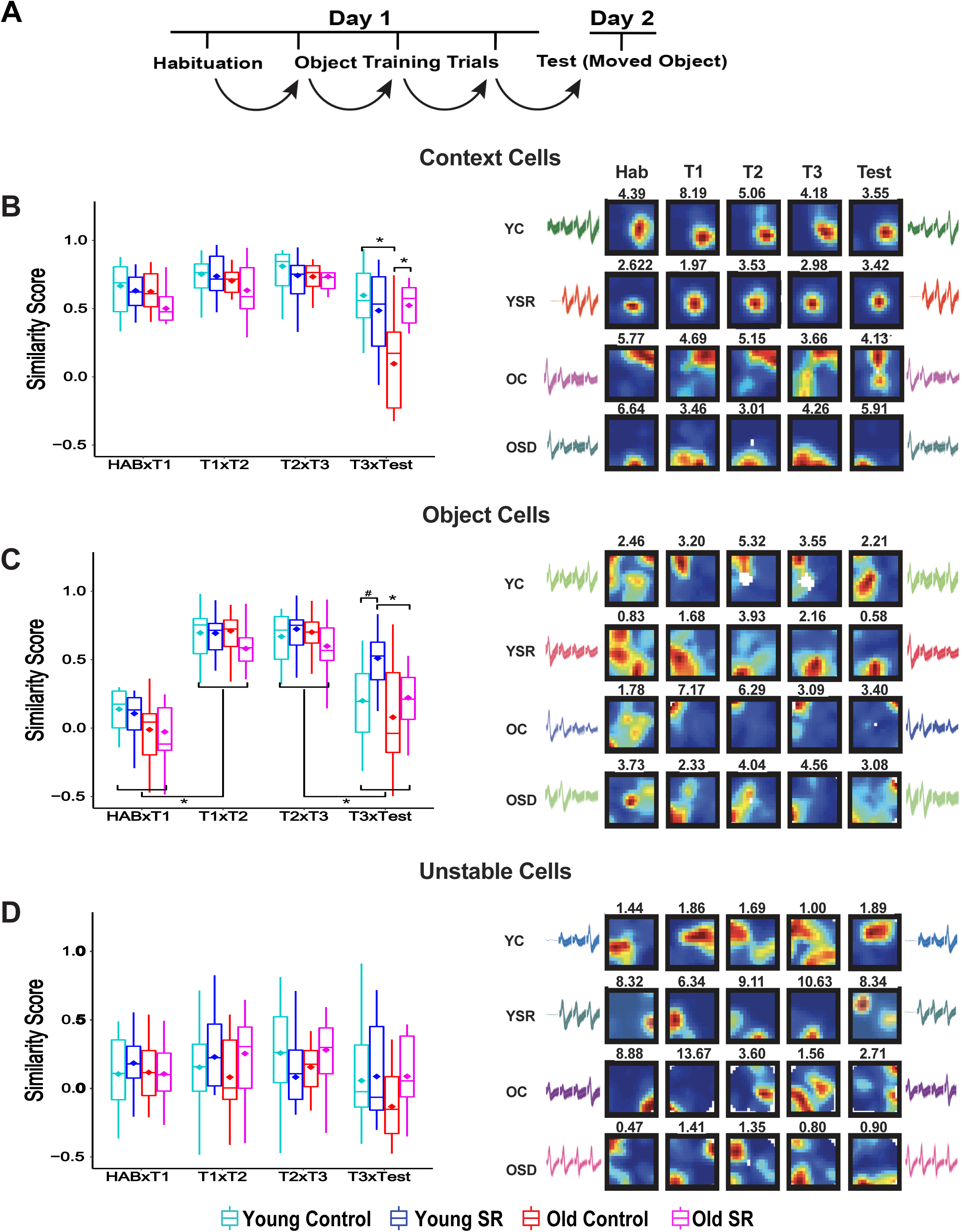
Cell-type remapping across sessions during performance in the OPR task. A) Schematic indicating how similarity scores were computed. B-D) Average global remapping of context (B), object (C), and unstable (D) cells across trials and corresponding examples of color-coded place cell rate maps and waveforms. Context cells are only unstable in the old control group, suggesting that poor performance in this group stems from inability to recall the context. Conversely, object cells, which by definition remap when the objects are introduced, display intermediate levels of stability in young controls and old SR, the groups that show successful performance in the OPR task. Conversely, poor OPR performance correlates with either too much object cell stability (young SR animals) or too much object cell instability (old controls). Blue indicates that the animal has visited the region but the cell has not fired, vivid colors indicate high neuronal activity. Number on top of each map represents peak firing rate (Hz) used to normalize the map colors. Waveform similarity indicates recording stability during the 24 hr period of training and testing. Context cells: Young controls (YC=23 cells), young SR (YSR=20 cells), old control (OC=8 cells), old SR (OSR=9 cells). Object cells: Young control (YC: 15 cells), young SR (YSR: 13 cells), old control (OC: 17 cells), old SR (OSR: 14 cells). Unstable cells: Young control (YC=22 cells), young SR (YSR=17 cells), old control (OC=11 cells), old SR (OSR=18 cells). Asterisks (*) represent significance using alpha=0.05. Number sign (#): indicates statistical trend. Statistical details in Supplemental Tables.

We also hypothesized that if animals noticed the change in the object configuration, object cells should display remapping during the test. However, if animals failed to notice the moved object, these cells should remain stable. Object cells displayed an effect of age [Fw(1)=18.71, p<0.0001] and trial [Fw(3)=24.30, p<0.0001] and an interaction between age and trial [Fw(1)=3.88, p<0.0001], as well as trend in the interaction between sleep condition and trial [Fw(3)=0.68, p<0.06]. Analysis of simple effects showed that all the groups displayed remapping between Hab and T1 (Zw=12.52, p<0.00001), a characteristic that served to categorize these cells, as well as between T3 and Test (Zw=7.31, p<0.0001). During the test, young SR animals displayed more stability than old SR mice [Tw(59.48)=4.34, p<0.0001], and a trend in comparison to young controls [Tw(50.19)=1.87, p=0.07]. This reflected that object cells from young SR were stable in comparison to the other groups. Successful memory performance in young controls and old SR correlated with intermediate levels of object cell remapping (**Figure 3C**). Finally, there were no stability differences in “unstable” cells across groups or trials [age: Fw(1)=0.35, p=0.56, sleep: Fw(1)=0.76, p=0.51, trial: Fw(3)=0.76, p=0.52, age vs. sleep vs. trial: Fw(3)=0.43, p<0.73; **Figure 3D**]. Together, these data indicate that the successful memory performance observed in young controls and old SR mice correlates with stable context cells and flexible object cells.

To evaluate if the remapping observed in young and old control animals reflected different mnemonic processes, the former being associated with memory updating and the latter reflecting typical long-term instability, we conducted an unmoved object control task in a subset of animals. Three young mice (30 cells) and four old mice (24 cells) were tested in a non-moved object control task. Animals were trained exactly as described in the OPR task; however, during the test, all objects remained in the same position (**Figure 4A**). We found that object and context cells were stable in young mice, but not in old mice. Unstable cells displayed similar remapping characteristics in both groups (Figure 4B-C). These data indicate that the remapping observed in young control mice during the moved object test reflects memory updating rather than passage of time and the high instability of object cells in old controls is inherent in this group.

**Figure 4.**
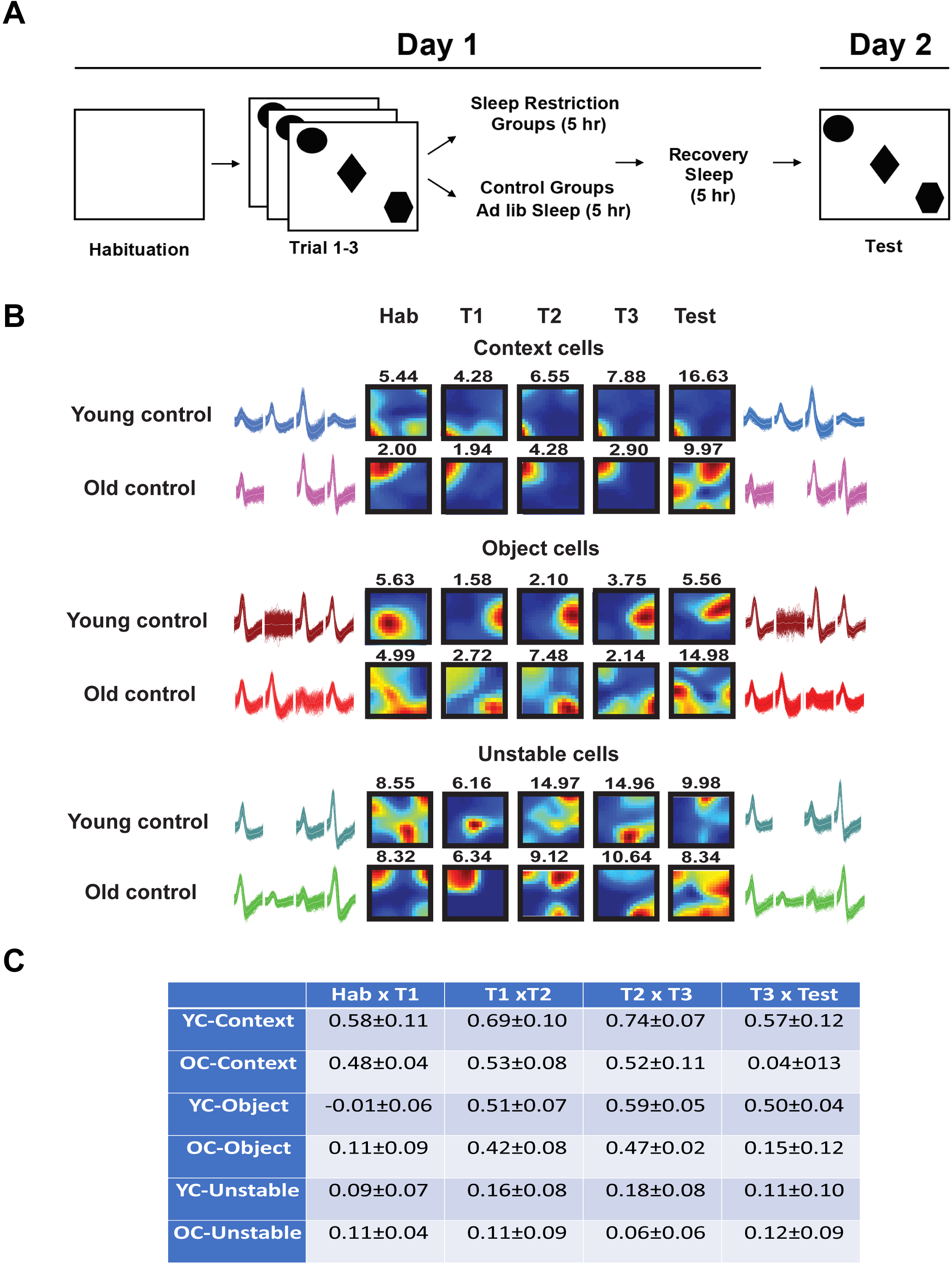
Place cell stability during unmoved object control task. A) Schematics of the task. Training was identical to that in the OPR task, but all objects remain in the same position during the test. B) Examples of context, object, and unstable cells and waveforms recorded from young and old control mice trained in the unmoved object control task. Context and object cells recorded in young mice are stable between the last training trial and the test, whereas all cell types are unstable in old mice. In the place cell rate maps, blue indicates that the animal has visited the region but the cell has not fired, vivid colors indicate high neuronal activity. Number on top of each map represents peak firing rate (Hz) used to normalize the data. Waveforms similarity indicates recording stability during the 24 hr period of training and testing. C) Average similarity scores for context, object, and unstable cells recorded from young and old mice during OPR unmoved object control task. Young: 3 mice, 30 cells, old: 4 mice, 24 cells.

### Sleep analysis: Bayes classifier validation

Sleep patterns from young an old animals were recorded in chambers that provided free access to food and water (**Figure 5A)**. To analyze all the wake/sleep periods we used a validated Bayes classifier to minimize inter-observer variability (Grigg-Damberger, 2012). This classifier that has been previoulsy tested in control, drug infused, aged, and transgenic mice (Rytkonen et al., 2011). To visualize the accuracy of the classifier in our data set, we color coded the EMG and EEG using the output of the Bayes classifier to determine that there were no significant errors in the scoring of Wake, NREM, and REM states (**Figure 5B**). Then, we partially re-validated this method comparing manual visual scoring with the output of Bayes classifier. A confusion matrix was created for each animal and an average matrix was generated by averaging values for each age group (4 young and 4 old mice, Figure 5C-D). Accuracy was calculated by adding the values in which algorithm and visual scoring correlated in Wake, NREM and REM divided by the total entries. The results indicated that the algorithm was very accurate for both young and old mice (**Figure 5E**). Then, we calculated sensitivity (true positive rate) and specificity (true negative rate) for both young and old animals to generate a receiver-operating curve (ROC, see method for description of calculations). The results indicated that the classifier generated outputs with high positive rates and minimal false positives (**Figure 5F**). Moreover, we compared the reliability of the classifier using 5% and 10% of manually scored data obtaining highly similar results (average inter-score correlation: 95 ± 0.54%, 3 mice).

**Figure 5.**
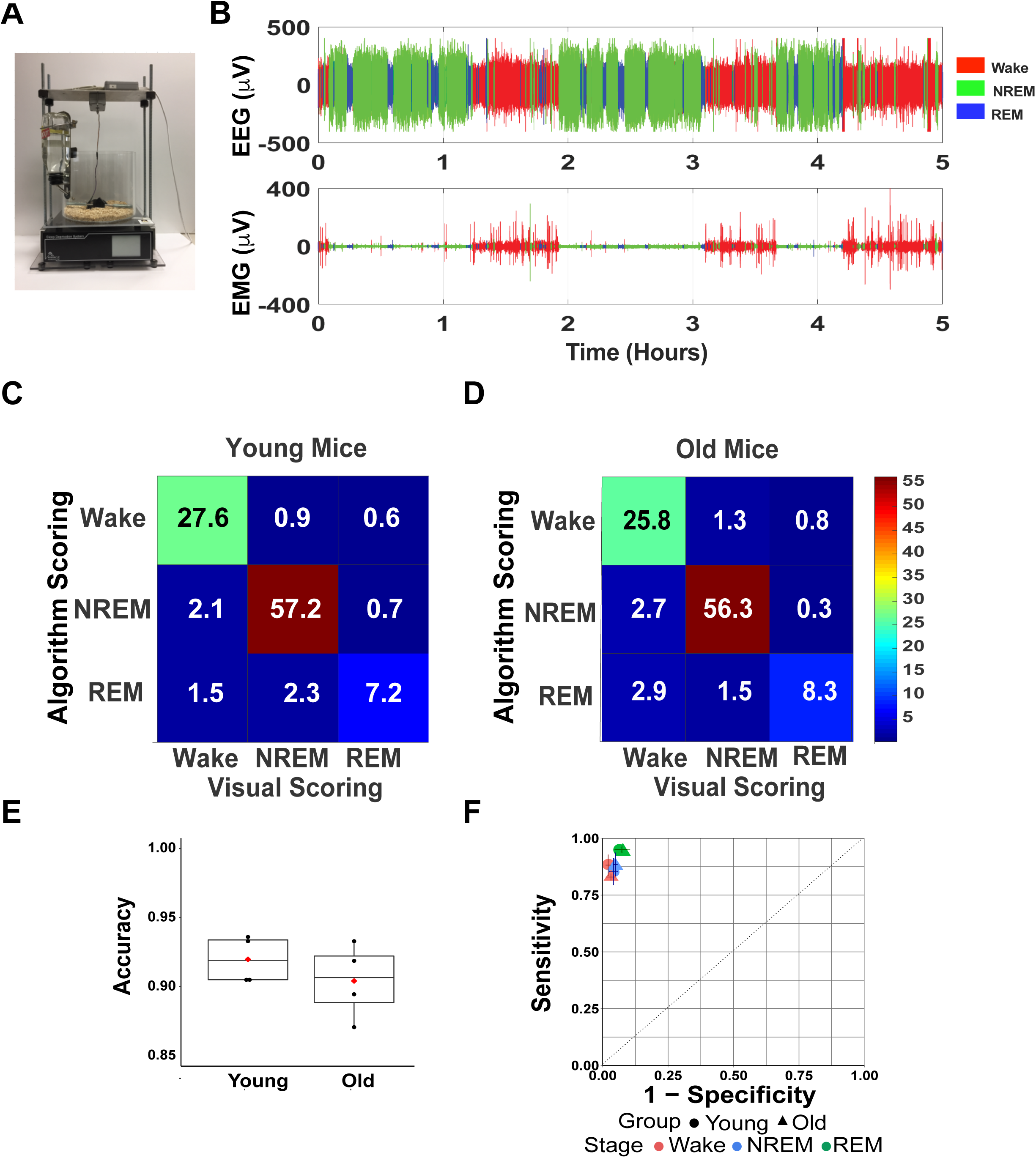
Bayes classifier validation. A) Photograph of the sleep restriction chamber. B) Representative 5 hr sleep recording showing electroencephalogram (EEG, top panel) and electromyograph (EMG, bottom panel) color-coded using the output of a Bayes classifier. The graph illustrates the accuracy of the classifier detecting Wake, NREM and REM periods. C-D) Average normalized confusion matrices for young (C) and old (D) mice. To determine the accuracy, sensitivity and specificity of the classifier, a confusion matrix was created by comparing visual and algorithm scored data in 4 young and 4 old mice. The results of this comparison were summed into a confusion matrix for each animal. Each entry of the confusion matrix was then divided by the total number of epochs and multiplied by 100 to turn the entries into percentages. Average (arithmetic mean) confusion matrices were then computed for each age group (young and old, respectively), which are shown in panel C. E) Accuracy of the Bayes classifier for young and old mice. F) Receiver operating characteristic (ROC) curve showing average values of sensitivity (true positive rates) against 1-specificity (false positive rates) for average values calculated in Wake, NREM, and REM in young and old mice. The high accuracy of the classifier is illustrated by the fact that all values are plotted in the upper left area, indicating high sensitivity and low fall-out errors.

### Old control mice display more fragmented NREM sleep patterns than young mice during post-learning ad libitum sleep

We analyzed sleep patterns in the control groups during the post-learning *ad libitum* sleep period. We found that the total percent time spent in Wake, NREM and REM was equivalent in young and old control animals [Wake: Tw(10.33)=0.91, p=0.39, NREM: Tw(10.47)=0.49, p=0.63, REM: Tw(10.28)=1.42, p=0.19, **Figure 6A**]. However, old animals displayed more NREM and REM bouts and a non-significant increase in Wake bouts than young controls [NREM: Tw(9.23)=2.84, p<0.02; REM: Tw(8.46)=2.97, p<0.02, Wake: Tw(6.75)=1.37, p=0.22; **Figure 6B**].

**Figure 6.**
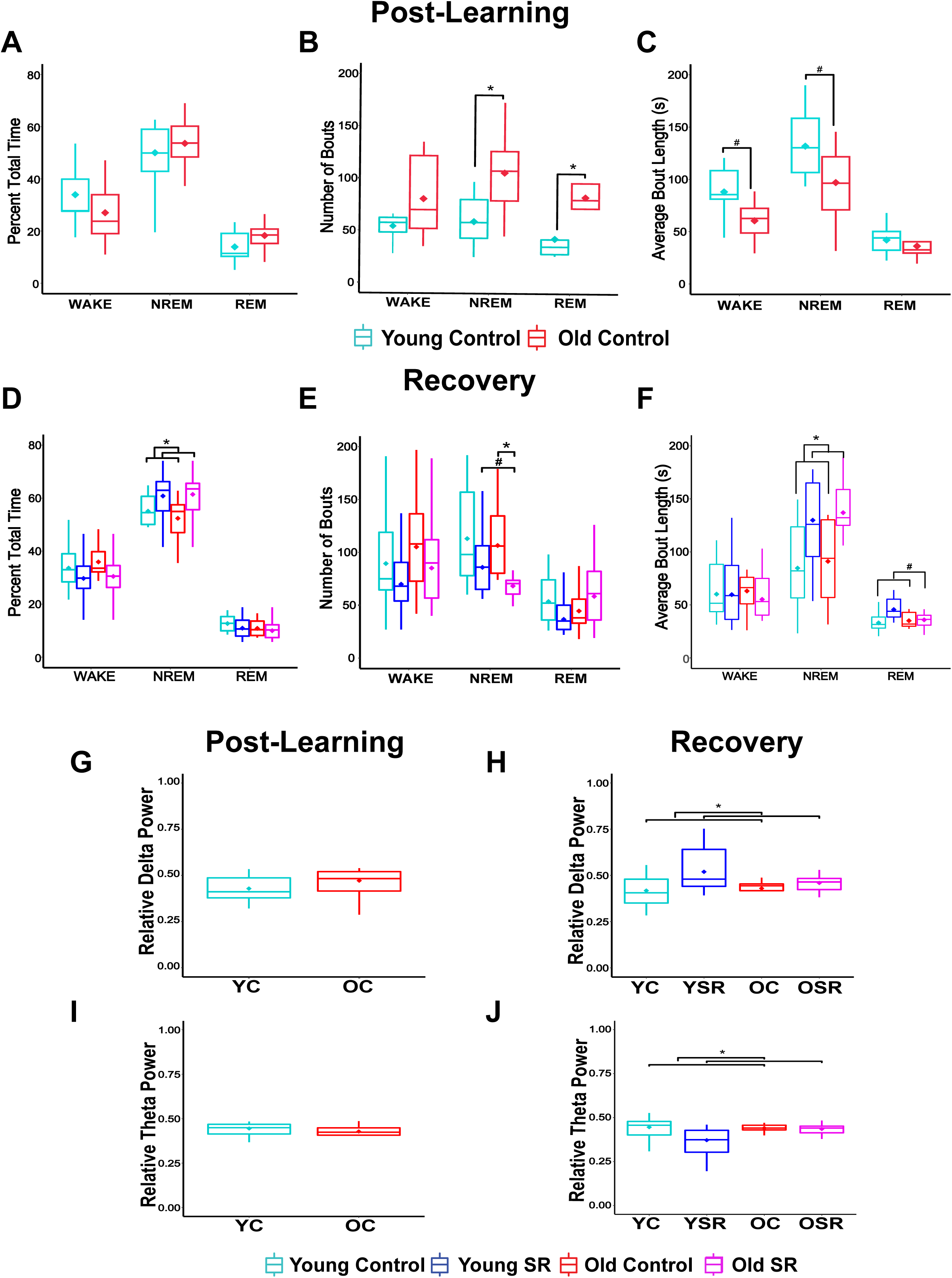
Sleep patterns and relative delta and theta power during post-training and recovery sleep periods in young and old mice. A-C) Percentage of total time (A), number of bouts (B), and bout length (C) in Wake, NREM, and REM during post-training sleep in young and old control mice. D-F) Percentage of total time (D), number of bouts (E), and bout length (F) in Wake, NREM and REM during recovery sleep in control and SR groups. G-I) Percentage of total time (G), number of bouts (H), and bout length (I) in Wake, NREM, and REM in young and old control animals during post-learning and recovery periods (within subject analysis). G-H) Relative delta power (RDP) in NREM during post-learning (G) and recovery (H). I-J) Relative theta power in REM during post-learning (I) and recovery (J). Relative delta power (RDP, 0.25-4 Hz), relative theta power (RTP, 4-10 Hz),. Young control (n=11), young SR (n=11), old control (n=8), old SR (n=10). Asterisks (*) represent significance using alpha=0.05. Number sign (#) indicates statistical trend. Statistical details in Supplemental Tables.

Additionally, old mice displayed a trend toward shorter Wake and NREM bout length than young controls [Wake: Tw(9.85)=1.92, p = 0.09; NREM: Tw(10.97)=2.09, p = 0.06], but no differences in REM bout length [REM bout length: Tw(10.54)=1.01, p=0.34, **Figure 6C]**. These data indicate that old animals display more fragmented NREM sleep than young mice, as previously observed (Pace-Schott and Spencer, 2015).

Next, we analyzed sleep patterns in the SR groups during the first 5 hr post-learning to corroborate that the animals remained awake during this period. On average SR mice were awake 97.5% of the time (young animals: 97.55±1.26%, old animals: 97.51±1.59%, data not shown), indicating that the sleep restriction procedure was effective.

### Old mice display increased NREM consolidation during recovery sleep following sleep restriction

Following the initial 5 hr post-training period, we recorded 5 additional hours of *ad libitum* sleep in the control and SR groups to evaluate the effects of SR on recovery sleep. Old and young animals did not display differences in total time spent in Wake or REM [Wake: age: Fw(1)=0.33, p=0.57, sleep: Fw(1)=3.02, P=0.10, age vs. sleep: Fw(1)=0.04, p=0.85, REM: age: Fw(1)=0.84,

p=0.36, sleep: Fw(1)=0.76, P=0.39, age vs. sleep: Fw(1)=0.08, p=0.68, **Figure 6D**]. However, there was an effect of sleep condition in NREM [Fw(1)=4.65, p<0.05], but no age or interaction effects [age: Fw(1)=0.09, p=0.77, age vs. sleep: Fw(1)=0.06, p=0.74], which reflected that both young and old SR animals displayed more time in NREM (**Figure 6D)**.

Next, we analyzed changes in sleep microstructure during Wake, NREM, and REM. There were no significant changes in Wake bout numbers or length [bout number: age: Fw(1)=1.23, p=0.28, sleep: Fw(1)=1.99, P=0.18, age vs. sleep: Fw(1)=0.05, p=0.82, bout length: age: Fw(1)=0.40, p=0.54, sleep: Fw(1)=0.02, P=0.90, age vs. sleep: Fw(1)=0.01, p=0.97, Figure 6E-F]. Conversely, there was an effect of sleep condition [Fw(1)=5.34, p<0.05] and an interaction between age and sleep condition in NREM bout number [Fw(1)=0.32, p<0.04]. Analysis of this interaction showed that old SR mice displayed fewer NREM bouts than old controls [Tw(5.66)=2.73, p<0.04], but there were no differences between the young groups [Tw(8.88)=1.04, p=0.33; **Figure 6E]**. Additionally, there was an effect of sleep condition in NREM bout length, reflecting that both young and old SR groups displayed longer NREM bouts than controls [Fw(1)=7.53, p<0.02, **Figure 6F**]. The observation that old SR mice displayed fewer numbers of NREM bouts of longer length than old controls indicated that NREM was consolidated in this group.

Finally, there was an interaction between age and sleep condition in the number of REM bouts [T_W_(1)=-0.36, p<0.05, **Figure 6E**] and trends in both the main effect of sleep condition [Fw(1)=4.19, p=0.054] and an interaction between age and sleep condition in REM bout length [Fw(1)= 0.34, p=0.059, **Figure 6F**]. These differences were driven by the fact that young-SR mice displayed less REM bouts of longer length, but these differences were modest and did not reach significance in *post hoc* simple tests [Number of bouts: YC vs. YSR: Tw(9.56)=1.41, p=0.19, OC vs. OSR: Tw(9.83)=0.84, p=0.37]. In summary, these data indicate that acute SR serves to consolidate NREM in old mice, only producing subtle sleep changes in young mice.

### Acute sleep restriction increases relative delta power (RDP) during NREM and decreases relative theta power (RTP) during REM in SR mice

Delta and theta oscillations have been associated with cognitive tasks requiring attention, spatial exploration, and memory (Harmony, 2013; Hasselmo and Stern, 2014; McKenzie and Buzsaki, 2016). Therefore, we first investigated if these oscillations were differentially altered during the sleep restriction period in young and old mice. To this end, we quantified relative delta (RDP) or theta (RTP) power during SR finding that there were no differences between the groups (RDP: Young-SR: 0.29±0.03, old-SR: 0.33±0.02; RTP: Young-SR: 0.44±0.03, Old-SR: 0.48±0.02, p>0.05; data not shown). These results indicated that the movement of the bar did not differentially affect the EEG in the experimental groups.

Delta power has also been shown to increase following sleep deprivation (Davis et al., 2011). However, this effect depends on the duration of wakefulness (Dispersyn et al., 2017; Halassa et al., 2009), animal housing conditions (Kaushal et al., 2012), and several other variables (Davis et al., 2011). To determine if the experimental conditions of this study affected delta power, we calculated RDP in our groups. There were no significant differences in RDP between young and old control mice during *ad lib* sleep following training [p>0.05, **Figure 6G**]. However, there was an effect of sleep condition during recovery (Fw=5.17, p<0.03; **Figure 6H**), reflecting an increase in RDP in the SR groups, as previously shown (Halassa et al., 2009).

Mendelson and Bergmann (1999) found that normalizing the delta power in NREM by the delta power in REM revealed age differences (Mendelson and Bergmann, 1999). Therefore, we also tested this normalization method in our data. We again found moderate increases in RDP in the SR conditions, but no significant interactions [age: Fw(1)=0.66, p=0.43, sleep: Fw(1)=2.56, p=0.13, interaction; Fw(1)=0.35, p=0.68, Young control: 2.89 ± 0.40; Young SR: 3.98 ± 1.30; old control: 2.89 ± 0.37; Old SR: 3.71 ± 0.27, data not shown].

Theta oscillations during REM have also been associated with memory encoding (Hasselmo, 2006; Hutchison and Rathore, 2015). Therefore, we examined RTP during REM when this oscillation is most prominent. There were no significant differences during the post-learning *ad lib* sleep period (p>0.05, **Figure 6I**). However, there was an effect of sleep condition during the recovery (Fw=4.46, p<0.04; **Figure 6J**), indicating that the SR groups displayed less RTP than the controls, but no interactions between age and sleep condition were observed (p>0.05). In summary, these data suggest that 5 hr of SR followed by 5 hr of recovery sleep do not result in age differences in RDP or RTP between young and old mice.

### Young control mice display more spindles of longer length than old control mice immediately following training

Spindles appear to play a role in memory consolidation by facilitating memory reactivation (Rasch and Born, 2013). To address if spindles played a role in the memory enhancement observed in young controls and old-SR animals, we quantified spindle characteristics in our groups using a highly accurate automated spindle detection method (Uygun et al., 2019). First, we performed a partial re-validation of the method obtaining 95% reliability (n=4; young: 97% overall correlation; old: 94% overall correlation). Errors included 2.5% of false positives and 2 % of false negatives. Then, we reproduced the method developed by Uygun et al. and color-coded the detected spindles in the filtered EEG to visually inspect the reliability of the code (**Figure 7A**), observing very accurate results.

**Figure 7.**
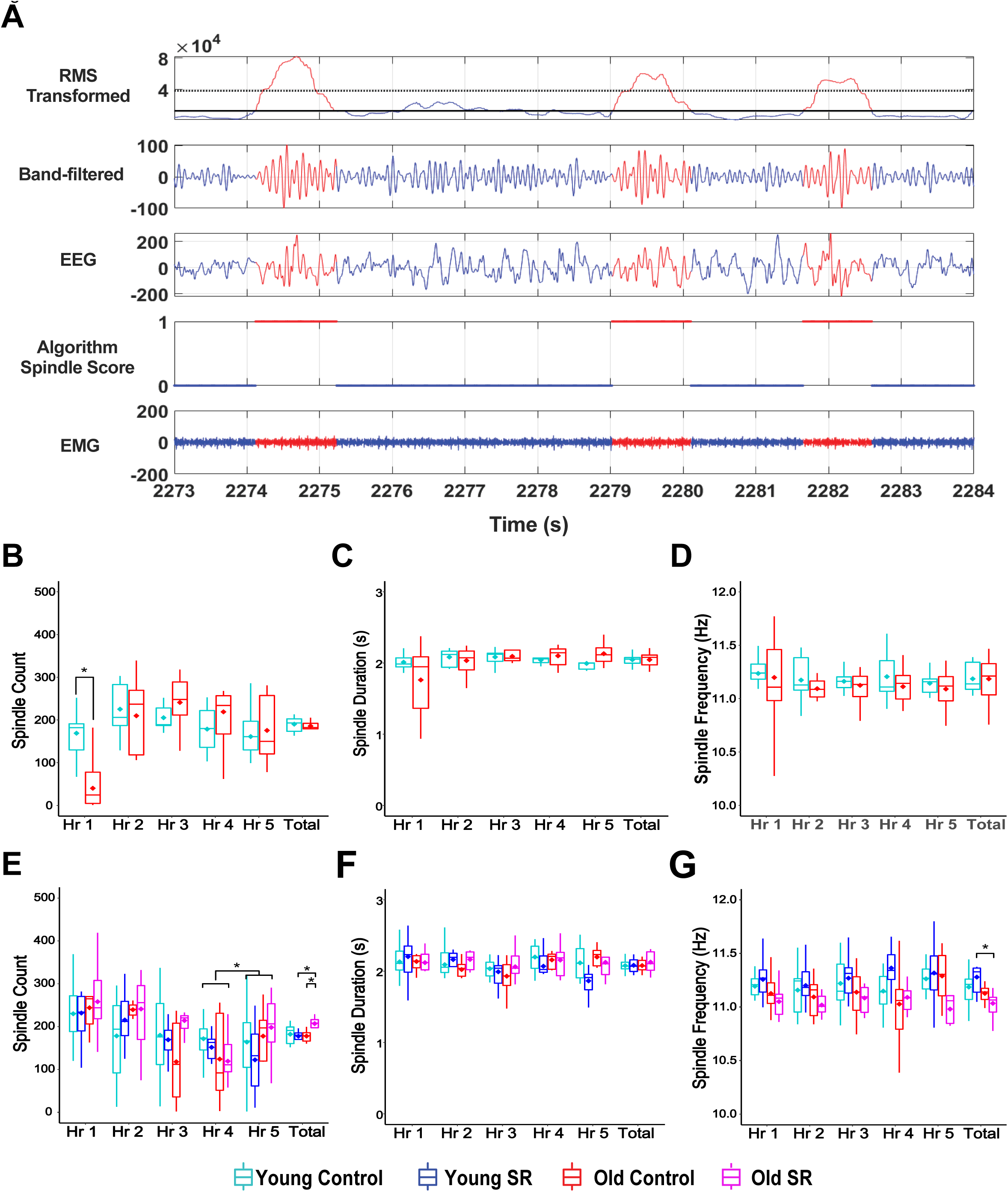
Spindle characteristics. A) Representation of the automated spindle detection method. Top panel shows the cubed root-mean squared (RMS) of the filtered EEG computed over 750 ms windows. The spindle detection method used a two-threshold approach. These values are derived from the mean cubed RMS transform value of all the NREM traces and included both a lower threshold and upper threshold. The second panel shows the band-pass filtered EEG, followed by the unfiltered EEG signal, and the results of the spindle detection. The bottom panel shows the corresponding EMG signal. Red marks the regions of the signal were spindles are detected, 1 indicates the presence and 0 the absence of spindles, respectively. B-D) Spindle counts (B), duration (C), and frequency (D) during the post-learning sleep period. B) Although there were no differences between the groups in total number of spindles, the segment analysis revealed that young control animals displayed more spindles than old control animals during the first hour post-learning. C-D) There were no differences in spindles duration (D) or frequency (D) during post-learning. E-G) Spindle numbers (E), duration (F), and frequency (G) during recovery. G) Old SR animals displayed more spindles than old controls during the recovery sleep period. The analysis during the 1 hr segments showed that old SR mice showed a trend toward more spindles across all segments, but the differences were only significant during the last segment (hour 5). F) There were no differences in spindles duration. G) Young SR mice displayed higher spindle frequency than old SR animals. Asterisks (*) represent significance using alpha=0.05. Statistical details in Supplemental Tables.

Using this method, we analyzed spindles during NREM periods. There were no significant differences in average number, frequency, or duration of spindles between young and old control mice during the *ad libitum* post-training sleep period [spindle count: Tw(10.99)=0.52, p=0.62, spindle duration: Tw(9)=0.004, p=0.99; spindle frequency: Tw(8.19)=0.01, p=0.99, Figures 7B-D]. Since the initial hours following training have been shown to be critical for memory consolidation in young animals (Bailey et al., 2004), we divided the post-training sleep period into consecutive 1 hour segments to investigate whether there were differences at distinct stages following learning. We found a significant main effect of hour segment and an interaction between age group and hour segment in spindle numbers [hour segment: Fw(4)=8.73, p<0.008; interaction: Fw(4)=4.35, p<0.05]. Analysis of simple effects indicated that young control animals displayed more spindles than old controls during the first hour post-learning [Tw(10.06)=4.18, p<0.002; **Figure 7B**]. No differences in spindle duration or frequency were observed in the hour segment analysis [spindle duration: age: Fw(1)=0.06, p=0.82, hour: Tw(4)=0.64, p=0.65, age vs. hour: Tw(4)=1.15, p=0.41, spindle frequency: age: Fw(1)=0.22, p=0.65, hour: Tw(4)=0.81, p=0.64, age vs. hour: Tw(4)=0.34, p=0.89, Figures 9C-D]. These results indicate that increased number of spindles during immediate post-learning sleep may facilitate memory consolidation in young mice.

### An acute session of sleep restriction increases total number of spindles in old animals during recovery

Previous research indicated that the time window for memory consolidation is much longer in old animals than young subjects (Schimanski and Barnes, 2010). These findings suggest that increases in spindle count occurring even several hours following learning may have a significant impact on memory consolidation in old animals. We examined spindle count, duration, and frequency during the recovery *ad libitum* sleep period following SR. We found a significant interaction between age group and sleep condition on spindle count [Fw(1)=1.00, p<0.02]. Analysis of single effects revealed that old SR mice displayed more spindles than old controls [Tw(8.85)=3.29, p<0.01, **Figure 7E**]. Although there were no differences in spindle duration during recovery [age: Fw(1)=0.23, p=0.64, sleep: Fw(1)=0.30, p0.59, age vs. sleep: Fw(1)=1.0, p=0.66, **Figure 7F**], there was an interaction between age and sleep condition on spindle frequency [Fw(1)=0.37, p<0.05]. This effect was driven by young SR animals displaying higher spindle frequency than old mice [Tw(10.29)=-3.73, p<0.004; **Figure 7G**]. This age difference parallels previous observations in humans following SR (Rosinvil et al., 2015).

We then examined if there were differences in spindle characteristics across the groups at different times during recovery by subdividing this period into 1 hr segments. We found a significant effect of hour segment on spindle count [Fw(4)=0.90, p<0.0003], and an interaction between age and sleep condition [Fw(1)=3.04, p<0.00001, **Figure 7E**]. Analysis of single effects indicated that the Old SR animals displayed a trend toward more spindles than young SR mice across all segments (Tw= -1.91, p=0.07), but the differences reached significance during the last segment (Zw=3.05, p<0.03, **Figure 7E**). There were no significant hour segment, sleep, or interaction differences in spindle duration or frequency [spindle duration: sleep: Fw(1)=0.01, p=0.97, hour: Fw(4)=0.37, p=0.83, age vs. sleep vs. hour: Fw(4)=0.08, p=0.99; spindle frequency: sleep: Fw(1)=0.01, p=0.96, hour: Fw(4)=0.17, p=0.96, age vs. sleep vs. hour: Fw(4)=0.09, p=0.99, Figures 7F-G]. Together, these results indicate that during recovery there is an increase in average number of spindles in old SR mice in comparison to old controls, which may serve to consolidate memory and facilitate performance of the task.

## Discussion

In this study, sleep restriction impaired OPR memory in young adult mice, but unexpectedly enhanced performance in old mice. The improved performance observed in old SR animals was accompanied by decreased NREM fragmentation and increased number of spindles during recovery sleep. Successful OPR in both young control and old SR mice correlated with stability of context cells representing the static aspects of the context and remapping of object cells coding the object configuration. Moreover, while both old control and young SR mice failed to recognize the displaced object, they exhibited different patterns of stability during the test session, indicating that performance deficits in these groups stemmed from distinct memory impairments. These data indicate that age-related cognitive deficits can be rescued by improving sleep architecture, which correlates with enhanced flexibility and stability of hippocampal representations.

The unexpected finding that OPR memory was enhanced in old SR mice suggests that SR followed by recovery sleep may have mnemonic advantages in old subjects. Sleep fragmentation has been associated with impaired memory consolidation (Sportiche et al., 2010; Tartar et al., 2006; Ward et al., 2009a; Ward et al., 2009b); thus, NREM consolidation may underlie the rescue of age-related cognitive deficits. Indeed, studies have found that enhancing NREM sleep via pharmacological interventions or SR can provide protective cognitive effects following stroke or traumatic brain injury (Cam et al., 2013; Martinez-Vargas et al., 2012; Morawska et al., 2016). Moreover, changes in NREM activity predict memory performance (Ognjanovski et al., 2014; Ognjanovski et al., 2017) and disruptions of hippocampal oscillations during NREM sleep disrupt memory consolidation (Ognjanovski et al., 2018). Our findings support these data by showing that NREM microstructure is important for memory consolidation.

Previous studies examining place cell activity during the OPR task have been conducted in young adult rats under normal *ad libitum* sleep conditions (Zheng et al., 2016). For example, Larkin et al. showed that hippocampal CA1 neurons exhibited changes in firing rate, but not in the cells’ preferred firing locations during the moved object test (Larkin et al., 2014). Similarly, we observed rate remapping between the last object trial and the test in young control and old SR mice. However, we also found that successful learning correlated with place field remapping in a subset of CA1 neurons during the test. The differences in stability between our observations and Larkin et al. may be related to the intrinsic stability differences between mice and rats (Kentros et al., 2004; Muzzio et al., 2009b), the different retention intervals used in these studies (5 min in Larkin et al. compared to ≥15 hr in our study), or the fact that we categorized distinct cell types, finding instability only in object/unstable cells.

It was previously thought that the stability of place cells was critical for learning (Kentros et al., 2004; Muzzio et al., 2009a). However, recent findings in mice found that hippocampal cells expressing cfos, an activity marker associated with the formation of memory engrams (Liu et al., 2014), are much more unstable than cells that do not express this early gene (Tanaka et al., 2018). This confirms that subsets of cells participating in memory processes are indeed unstable. Interestingly, unstable subpopulations coexist with stable ones, potentially having distinct mnemonic functions. These observations are in line with our finding that the instability of object cells is important for memory updating, while stable context cells may serve a distinct memory function.

The general consensus from other studies using the OPR task in young animals is that immediate post-training sleep is critical for memory. Performance is optimal when the retention interval occurs during the inactive phase of the light/dark cycle and rats are allowed to sleep (Binder et al., 2012), but post-training SR has consistently shown to impair OPR memory in young adult rats (Sawangjit et al., 2018) and mice (Havekes et al., 2014; Prince et al., 2014). Our behavioral results in young mice are in agreement with these findings. However, although previous behavioral deficits have been attributed to complete disruption of memory consolidation, our results suggest otherwise. The long-term stability of context cells in young SR mice suggested that contextual representations were retrieved correctly, whereas the stability of object cells suggested that these mice failed to recognize changes in object configuration. Furthermore, if the object is not moved during the test, young control mice do not show object cell remapping, suggesting that the object cells’ shift in firing location reflects memory updating. The synaptic homeostasis hypothesis proposed by Tononi and colleagues posits that rather than actively strengthening memories, sleep may instead serve to downscale synapses in order to allow further memory acquisition (Tononi and Cirelli, 2006). Our results suggest that in young adult animals, SR may interfere with this process by making hippocampus-dependent memory more rigid and less flexible.

Similarly to Wimmer et al., (2012), we observe OPR performance deficits in old control mice. Critically, we demonstrate that the performance deficits observed in old mice are different from those found in young SR mice. In old mice, context cells show long-term instability, remapping between the training and test session, whereas in young SR mice context representations are stable. Moreover, instability of both context and object cells is also observed during the unmoved object control in old mice, suggesting that this is a characteristic of hippocampal representations in this group. Indeed place cell instability has been previously reported in old animals (Barnes et al., 1997), along with impairments in several spatial tasks (Rosenzweig and Barnes, 2003). Therefore, it is possible that the sleep fragmentation observed in old control mice contributes to impair consolidation of the static aspects of the environment.

It is well established that in young animals, the initial hours after training are critical for initiating transcriptional events that lead to the translation of new proteins important for memory consolidation (Bailey et al., 2004). We demonstrate that delaying the SR to begin 5 hours after training does not affect performance in young animals, as previously shown (Palchykova et al., 2006). Furthermore, young controls display more spindles, which are important for memory consolidation (Fernandez and Luthi, 2020), during the initial hour of post-training sleep. Interestingly, converging lines of evidence indicate that the time window for protein-synthesis dependent memory consolidation is extended in old animals (Schimanski and Barnes, 2010). Moreover, sleep loss inhibits translational processes in young mice, but not old ones (Naidoo et al., 2008). These observations suggest that NREM consolidation and increased spindle numbers during recovery sleep beginning 5 hours after learning may have beneficial effects in old mice because the changes occur when the window for memory consolidation is still open. However, the beneficial effects of SR on memory in old animals disappear when sleep loss happens 5 hr after training. This suggests that although the time window for consolidation is extended, it still has boundaries. Alternatively, delayed SR may not be beneficial because it pushes recovery sleep to the beginning of the dark cycle, when rodents are normally active, making the process more ineffective.

Spindles have been associated with memory encoding (Antony et al., 2019; Ulrich, 2016) and alterations in spindle count and properties are good predictors of age-related cognitive decline (Taillard et al., 2019). This likely happens because thalamocortical spindles lead to proliferation of hippocampal sharp waves (Clemens et al., 2006; Clemens et al., 2011; Isomura et al., 2006; Nishida and Walker, 2007; Peyrache et al., 2011; Siapas and Wilson, 1998; Sirota et al., 2003). Spindle oscillations coordinate activity of several cortical and subcortical regions in up and down membrane potential states (Isomura et al., 2006). In the hippocampus, the up states coincide with increased activity in CA1 and the dentate gyrus, as well as more sharp wave ripples events (Isomura et al., 2006). Since there is replay of wake activity in a compressed format during hippocampal ripples (Buzsaki, 2015; Joo and Frank, 2018), more spindles are likely to enhance memory. Our data support these findings by showing that increases in number of spindles even 5 hr post learning are sufficient to ameliorate age-related memory deficits in old animals.

In our experiments, SR produced a mild increase in RDP in SR animals and no age differences. Other studies using similar normalization methods reported age-dependent increases in RDP (Mendelson and Bergmann, 1999; Panagiotou et al., 2017). These discrepancies may reflect differences in procedures, including length of SR (48 hr vs. 5 hr) or recovery period (24 hr vs. 5 hr). Moreover, prolonged recovery recordings allowed identification of age differences at specific times of the sleep cycle. Old rats only displayed higher RDP than young ones during the first part of the dark cycle (Mendelson and Bergmann, 1999). We evaluated RDP during the second half of the dark cycle to train and SR animals in the first half. Therefore, our results are not in disagreement with previous reports but rather highlight the effects of distinct analysis times and/or procedures.

In summary, our findings contribute to a better understanding of the effects of sleep quality on memory and hippocampal representations, and have important potential clinical implications for rescuing age-related cognitive deficits.

## STAR Methods

### Subjects

Young (8-24 weeks old) and aged (54-72 weeks old) adult male C57BL/6J mice (Jackson Laboratory, Bar Harbor, ME) were housed individually on a 12-hour light/dark cycle and allowed access to food and water *ad libitum*. Animal living conditions were consistent with the standard required by the Association for Assessment and Accreditation of Laboratory Animal Care (AAALAC). All experiments were approved by the Institution of Animal Care and Use Committee of the University of Texas at San Antonio and were carried out in accordance with NIH guidelines.

### Surgery

For sleep recordings, prefabricated 2 EEG and 1 electromyograph (EMG) channel headmounts (Pinnacle Technology) were implanted [from Bregma (in mm): frontal leads: AP: +3.2, ML: ±1., and parietal leads: AP: -1.8, ML: ±1.2] and secured with cyanoacrylate and dental cement. Two EMG leads were placed under the nuchal musculature and affixed with VetBond. For place cell recordings, animals were implanted with custom made drivable six-tetrode microdrives made in our laboratory using Neuralynx connectors (EIB-36-narrow). The microdrives were affixed to the skull with cyanoacrylate and dental cement, with recording electrodes placed directly above the dorsal hippocampus [from Bregma (in mm): AP, -1.7; ML, -1.6; form dura; DV, -1.0]. A ground wire was connected to a screw placed on the contralateral side of the skull. Tetrodes were made with 0.25 μm insulated wires (California Fine Wire, Grover beach, CA) with impedance values ranging 300-600 KΩ at 1 kHz (1-1.5 KΩ before plating). Sleep patterns were not analyzed in mice implanted with tetrodes because the EMG lead made the tetrode implants unstable and noisy. Animals underwent at least one week of recovery prior to recordings.

### Sleep Restriction (SR)

An automated SR cylindrical apparatus (Pinnacle Technology, Lawrence, KS) containing a bar spanning the enclosure was used for all SR procedures. Animals were individually housed in the apparatus for at least 24 hours prior to the beginning of the experiments with fresh bedding, food, and water, and were returned to the apparatus in between trials. To induce SR, the bar was rotated continuously by a motor at approximately 3 rpm with random reversals in rotational direction to prevent subjects from acquiring brief sleep periods through possible adaptation to the pattern of rotation. All SR procedures followed the object exposures, and therefore, were started within 4 hrs of the start of the light cycle.

### Experimental Design and Statistical Analysis Behavioral Training

For all animals, behavioral procedures (e.g., exposure to the objects on day 1 and memory testing on day 2) were conducted during the first 4 hr of the light cycle (ZT 0-4). The object/place recognition task (OPR) was conducted in a square context (35cm x 35cm) with visual features on each wall for orientation (Figure 1A). Everyday items (glass beer bottle, metal soda can, and plastic juice bottle) were used as objects after pilot testing determined mice showed roughly equal preference for all items on average. On day 1, animals were habituated to the empty context for a 6 min habituation trial (Hab). After the habituation, the 3 objects were arranged along one of the diagonal axes of the context and three 6-min object exploration trials (T1-3) were conducted.

During the inter-trial interval (2 min), the animals were placed in their home cage and the context and objects were wiped down with 70% ethanol. Immediately following the third object trial (T3), animals in the experimental groups were housed in the experimental chamber and were sleep restricted for 5 hr, whereas controls were housed in the same chamber but allowed to sleep. On day 2, one object was moved from its original location to an adjacent corner and mice were tested for 6 min in the context with the moved object (Test). In the unmoved object control condition, animals were trained as described for the traditional OPR and allowed to sleep *ad libitum* following training, but all objects remained in their original locations during the test. In the delayed SR procedure, animals were also trained as described above and allowed to sleep *ad libitum* for 5 hr following training before undergoing the delayed 5 hr sleep restriction, followed by 5 hr of recovery sleep. In all conditions object positions were counterbalanced across trials.

### Behavioral Analysis

All object exploration trials were video recorded using Limelight (Actimetrics, Wilmette, IL) and analyzed offline by researchers blind to the group condition. All instances when an animal was oriented toward and touching an object with nose, vibrissae, and/or forelegs within 0-3 cm of the object were recorded as “object exploration”; contacting an object while passing or oriented away were not considered. Animals with an average object exploration time less than 10 s on any trial were excluded from analysis (2 mice were excluded). Behavioral data from animals used for sleep analysis was combined with data from animals used for place cell recordings.

Object preference was calculated as the percentage of time spent exploring the moved object in the test session relative to total object exploration time during testing minus the relative time exploring the same object during training, as previously described (Oliveira et al., 2010). This method estimated object preference taking into account any potential bias that animals may have had during training. Specifically, the percentage change in preference was calculated using the following formula:

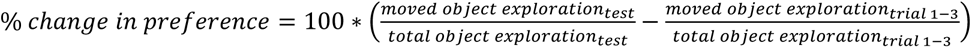

A larger percentage change indicates greater preference for the moved object during the test session, while lower values indicate little change in preference from day 1 to day 2, after the object is displaced. To rule out novelty effects, we repeated the same analysis excluding trial 1 when the objects were first introduced.

To calculate preference for the unmoved objects we used the following formula:

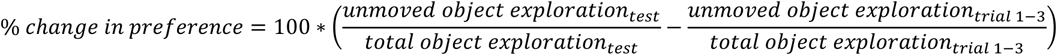

### Plasma corticosterone measures

Blood collection was conducted during the second quarter of the light cycle between 2-4 PM for all animals. Young and old animals were randomly assigned into control and experimental conditions between 8-10 AM. The experimental animals were sleep restricted for 5 hours, whereas control mice were allowed to sleep *ad libitum* in the experimental chamber. Following this period, all animals were deeply anesthetized with a mixture of ketamine and xylazine (100/10 mg/kg) and blood samples were extracted by cardiac puncture. Plasma was frozen and sent to Penn Diabetes Center Radioimmunoassays and Biomarkers Core for analysis (University of Pennsylvania, Philadelphia, PA).

### Place Cell Recordings and Analysis

Beginning one week after surgery, neural activity was screened daily in an environment different from the context used for experiments, advancing the electrode bundle 15-20 μm per day until pyramidal cells could be identified by their characteristic firing patterns (Ranck, 1973). Lowering the electrodes in small steps minimizes electrode drift and ensures recording stability for several days (Muzzio et al., 2009b; Wang et al., 2012). Moreover, all animals yielding unit data remained connected to the recording setup via a commutator throughout the experiment to further minimize the possibility of electrode drift during plugging/unplugging. Long-term recordings were considered stable when cells had the same cluster boundaries over two sessions (at least 24 hr apart), and the waveforms obtained from all four wires of a tetrode were identical. Experiments began only when recordings were stable for 24 hr. Animal position and electrophysiological data were recorded using Cheetah Data Acquisition system (Neuralynx, Bozeman, MN), as previously described (Wang et al., 2012; Wang et al., 2015). All tetrode implanted animals were trained in the OPR task as mentioned in the behavioral section. One subset of implanted animals was trained in the same manner but the objects remained unmoved during the test session. This control was added to determine if the observed changes in stability in some groups were due to memory updating or passage of time.

Units were isolated using MClust software (developed by A. David Redish, University of Minnesota) and accepted for analysis only if they formed isolated clusters with clear Gaussian ellipses and minimal overlap with surrounding cells and noise. All cells were inspected to rule out the presence of events during the 2 ms refractory period. Place field maps were generated using custom Matlab code as previously described (Keinath et al., 2014; Wang et al., 2012; Wang et al., 2015). Briefly, the arena was first divided into a 20×20 pixel grid and an activity map (the total number of spikes in each pixel), and a sampling map (the total amount of time spent in each pixel) were computed. Both maps were then smoothed with a 3 cm Gaussian kernel. The smoothed activity map was then divided by the smoothed sampling map, which yielded the place field map. Any location sampled for less than 1 s was considered un-sampled. Only periods of movement were included in the analysis (minimum walking speed: 2 cm/s). Cells that fired less than 25 spikes during movement or displayed peak firing frequencies below 1 Hz before smoothing were excluded from analysis. Firing rate patterns were characterized by computing the overall mean (MFR: total number of spikes divided by time spent in the arena), peak firing rate (PFR: maximum rate value), and out of field firing rate (OFFR: spikes occurring outside areas defined as place fields). Place fields were defined as any set of at least 9 contiguous pixels in which the average firing rate was at least 20% of the peak firing rate (Rowland et al., 2011). If a cell yielded multiple place fields, the sum of all fields was taken as the place field size. Rate remapping was calculated as the absolute difference between the peak firing rate of individual cells on consecutive trials. The spatial information content, a parameter that estimates how well the firing pattern of a given cell predicts the location of the animal, was computed as previously described (Skaggs et al., 1993) using the following formula IC=Σpi(Ri/R)log(Ri/R),where pi is the probability of occupying location i, Ri is the firing rate at location i, and R is the overall mean firing rate.

Place field stability was assessed by calculating pixel-to-pixel cross-correlations between maps. The generated Pearson R correlation value reflected the degree of map similarity across trials for all cells. Overall global remapping was estimated by averaging the Pearson r correlation values across cells and animals in each condition. Additionally, cell types were classified into three categories depending on whether they remapped in the presence of the objects (object cells), remained stable throughout training (context cells), or displayed both short- and long-term instability (unstable cells), with stability defined as a correlation value above 0.35, a threshold previously used in mice (Muzzio et al., 2009b; Wang et al., 2012).

### Verification of electrode placement

Tetrode placements were verified after completion of the experiments by passing a small current (0.1 mA) for 5 s through the tetrodes that yielded data in anesthetized animals. The brains were removed and fixed in 10% formalin containing 3% potassium ferrocyanide for 24 hr. The tissue was cryosectioned and stained using standard histological procedures (Powers and Clark, 1955).

### Sleep State Analysis

EEG/EMG signals were recorded for 10 hours following training on day 1 (Pinnacle Technology, Lawrence, KS). The headmounts were attached to a preamplifier for first stage amplification (100x) and initial high-pass filtering (0.5 Hz for EEG and 10 Hz for EMG). All signals were then sampled at 400 Hz and digitized (Sirenia Acquisition software, Pinnacle Technology). Animals with excessive noise in any channels (>10% of epochs classified as artifact) were discarded from analysis (2 animals were excluded from the post-training session due to noise and 2 animals were discarded from all sessions due to extremely noisy EEG and EMG).

Sleep recordings were divided into 4 s epochs. 5% of epochs were randomly selected for manual scoring with Sirenia Sleep analysis software using EMG power and EEG amplitude and frequency to categorize an epoch as a REM, NREM, or Wake states. Scoring randomly selected epochs produces more reliable outputs than continuous manual scoring of epochs (Rytkonen et al., 2011). The partially scored EEG/EMG files were then exported to MATLAB and the remaining epochs were analyzed using a naïve Bayesian classifier, a highly accurate method that has been shown to produce inter-rater agreements of 92% (Rytkonen et al., 2011).

### Classifier Validation

Complete visually scored data were compared with the output of the Bayes classifier to re- validate the method in a subset of animals. Additionally, outputs obtained with 5% and 10% manually scored inputs were compared to determine the effectiveness of the classifier using different amounts of input data in subsets of animals. To determine the accuracy, sensitivity and specificity of the classifier, a confusion matrix was created for each mouse visually scored. To this end, an experienced researcher visually scored 100% of the 4-second epochs comprising 5 hours of EEG and EMG activity. The researcher then selected only 5% of the epochs of the same recording. The 5% of visually scored epochs and the EEG and EMG recording were fed into the Bayes classifier to generated 5 hr of algorithm scored data. The scores from the researcher and classifier were then compared per epoch and added into the confusion matrix (y axis represents algorithm scoring and x axis visual scoring). Each entry of the confusion matrix was then divided by the total number of epochs and multiplied by 100 to turn the entries into percentages. An average (arithmetic mean) confusion matrix was then computed for each age group (young and old, respectively).

Confusion matrix measures

a, e, and I (shown in gray in the diagram below) are the boxes that represent values that algorithm and visual scoring coincided. Accuracy was calculated as follows:

Accuracy = (a + e + i) / (a + b + c + d + e + f + g + h + i)

Wake Sensitivity (True Positive) = a / (a + d + g)

Wake Specificity (True Negative) = (e + f + h + i) / (b + c + e + f + h + i)

NREM Sensitivity (True Positive) = e / (b + e + h)

NREM Specificity (True Negative) = (a + b + g + i) / (a + d + g + c + f + i)

REM Sensitivity (True Positive) = i / (c + f + i)

REM Specificity (True Negative) = (a + b + d + e) / (a + b + d + e + g + h)

Where the entries of a confusion matrix are denoted as follows:

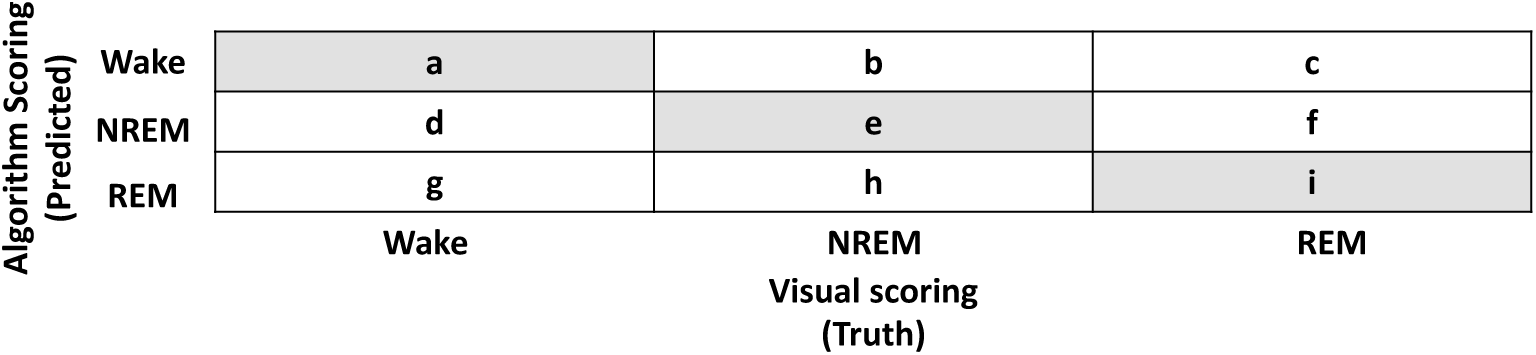

### Relative band power

The average power spectra were calculated use Welch’s power spectra method (MATLAB pwelch function), and normalized by total power as described previously (Mendelson and Bergmann, 1999). The relative band power (for a given state) was found by numerically integrating the power spectra over the given band’s frequency range using the MATLAB’s trapz function divided by the total power. A second normalization method was also used to evaluate relative delta power (RDP): Each 4 s segment of the scored EEG was multiplied by a 4 s long Hanning window. Then, a Fast Fourier transform (FFT) was calculated. The magnitudes of all the FFTs for any given state (Wake, NREM, or REM) were averaged to produce one frequency spectrum for each state. Total delta power was computed as described above using the trapezoid method. Relative power was calculated as the delta band power of NREM divided by the delta band power in REM (Mendelson and Bergmann, 1999). Delta frequency: 0.25-4 Hz, Theta frequency: 4-10 Hz.

### Spindle detention

Spindle detection during NREM was computed using a validated automated system for rapid and reliable detection of spindles using mouse EEG. This method eliminates observer bias and allows quantification of sleep parameters including count, duration, and frequency as well as rapid quantification during selective sleep segments (code generously provided by Dr. David Uygun). Briefly, the raw EEG signal is first bandpass filtered using the *designfilt* MATLAB function, which is then applied to filter the data using *the filtfilt* function with the following parameters: First stopband frequency = 3 Hz, first passband frequency = 10 Hz, second passband frequency 15 Hz, second stopband frequency = 22 Hz, with stopband attenuation levels of 24 dB. Once the EEG was filtered, the root-mean squared of the signal is calculated using a 750 ms window to smooth the data. This window was selected to minimize erroneous detection of artifacts. The RMS values were then cubed to enhance separation of signal and noise. Finally, spindles were detected using a two threshold approach that sets values from the mean cubed RMS transform of the NREM state. The lower threshold was calculated as 1.2 x mean cubed RMS and the upper threshold as 3.5 x mean cubed RMS (Uygun et al., 2019).

### Statistics

The results obtained through experiments are summarized in a Box-Plot graph according to robust statistics. Since the experimental data had irregular variability, e.g., some data did not fulfill the homoscedasticity assumption, we opted for robust statistical analysis, in order to protect the findings against such problems (Wilcox, 2012).

For the highest order design, the four groups of mice were arranged in a 2 Age (Young vs. Old) x 2 (Control vs. SR) Between Factorial Design, to analyze the independent effect of Age group or Sleep condition, compared to the combined effect (or interaction) between the two manipulations. In all analyses, we first tested Age by Sleep interactions. If there was a significant interaction, we tested for simple effects of Age on each individual Sleep condition, and the simple Effects of Sleep on each individual Age group, all this through an analysis of contrasts. When there was no significant interaction, the main effects analysis did not require post hoc tests since this was a 2×2 design. In some cases the design required 1-way analysis (equivalent to t-test since the variable had only two levels), while in other cases the design was more complex, requiring a 3-way type, e.g., 2×2 basic manipulation as well as the temporal trial (moment) manipulation as a factor of repeated measures.

Robust analyses of variance on trimmed means (Fw statistic) were conducted by using the method described by Wilcox (Wilcox, 2012), based on the generalization of Algina and Olejnik from the Johansen matrix algebra of factorial designs (Algina and Olejnik, 1984; Johansen, 1980). This statistic is distributed according to a Chi-squared, whose degrees of freedom (df) correspond to the number of contrasts necessary to decompose the variability of the effect, as is the case in the numerator of parametric F ANOVA (e.g. df=1 for main effects of Age, or Sleep Condition, and df=1*1 or 1, for interaction of both). Robust contrasts were conducted by a t-test trimmed variant (T_W_ on between, and Z_W_ on within factors), based on Yuen-Welch sequentially rejective post hoc test with Rom’s sequentially rejective method to control Type I error for multiple t-tests (see (Wilcox, 2012), for details). Specifically, this robust t-test is estimated from the squared standard error of trimmed means, calculated using the Winsorized variance. In the case of T_W_, the degrees of freedom are estimated with a logic similar to that of the t-student for samples with variances uneven, but replacing the standard error of the mean by the robust standard error of trimmed means (for details, see, Wilcox, 2012). In contrast, the Zw test does not have degrees of freedom associated with the test statistic since its value is based on a single pool of critical z values from a normal distribution, regardless of sample size. In order to gain sensitivity, given the complexity of some of the designs (e.g., 3-way mixed) and the presence of outliers, the probabilities associated with the Trimmed ANOVA were re-estimated using specific tests of first-order interactions (Patel and Hoel, 1973) and Bootstrap-t methods (Wilcox, 2012) in the case of second-order interactions. On the other hand, in the designs that included some variable of repeated measures, the basic test of contrasts was re-estimated using Percentile Bootstrap (Wilcox, 2012). All the functions that allow the robust ANOVA to be carried out according to all the types of designs used in the research are specified below. Finally, in all these analytical steps, we added the estimated effect size from Robust Wilcox “Explanatory Measures” (*Ef. Size* on Extended Data Tables). Values of .15, .35, and .50 correspond to the three bands of interpretation of the effect size as small, moderate, and large, respectively, from Wilcox robust interpolation of classical .2, .5, and .8 Cohen bands. Statistical significance was set at alpha 0.05. Data are graphed using box plots, plus the trimmed mean (trimmed = 0.2).

All statistical analyses were performed using the free-GNU R software, version 4.0.0 (Team, 2020) with *parallel*, *data.table*, *Hmisc*, and *ggplot2* libraries, as general libraries, and Wilcox’ *Rallfun-v37.txt* library (https://dornsife.usc.edu/labs/rwilcox/software/) for specific robust ANOVAs. Ask for the functions *bbwtrim*, and *bbwtrimbt* for 3-way Mixed designs [e.g. 2 Age (Young, Old) x 2 Sleep (Control, SR) x 5xS Trials (HAB, T1, T2, T3, Test)]; *t2way*, *t2waybt*, *rimul*, *bdm2way*, *ESmainMCP*, and *esImcp* for 2-way Between designs [e.g. 2 Age (Young,Old) x 2 Sleep (Control, SR)]; *bwtrim*, *bwtrimbt*, *ESmainMCP*, and *esImcp* for 2-way Mixed designs [e.g. 2 (Young, Old) x 5xS Hours (Hour 1 to Hour 5)]; *t1way* for 1-way Between design [e.g. 2 Age (Young Control, Old Control)]; *yuen*, *yuenv2*, and *lincon* for contrasts (or t-tests) with independent samples, and *yuendv2*, *rmmcp*, and *rmmismcp* for contrasts (or t-tests) for dependent samples.

## Code accessibility

All code and analysis tools will be available upon request.

## Data accessibility

All data will be uploaded on Mendeley

## Acknowledgments

This works has been funded by NSF (NSF/IOS 1924732 to IAM), NIH (R01 MH123260-01 to IAM, F31MH105161 to RKY and RISE GMO60655 to MRL, and IK2 BX004905 to DSU). We thank Sriharshini Muthukumar for running some behavioral experiments. In particular, we thank David S. Uygun, Fumi Katsumi, Robert E. Strecker and James M. McNally, Department of Psychiatry, VA Boston Healthcare System and Harvard Medical School, for providing a validated automated spindle detection code.

---

**Supplemental Figure 1.**
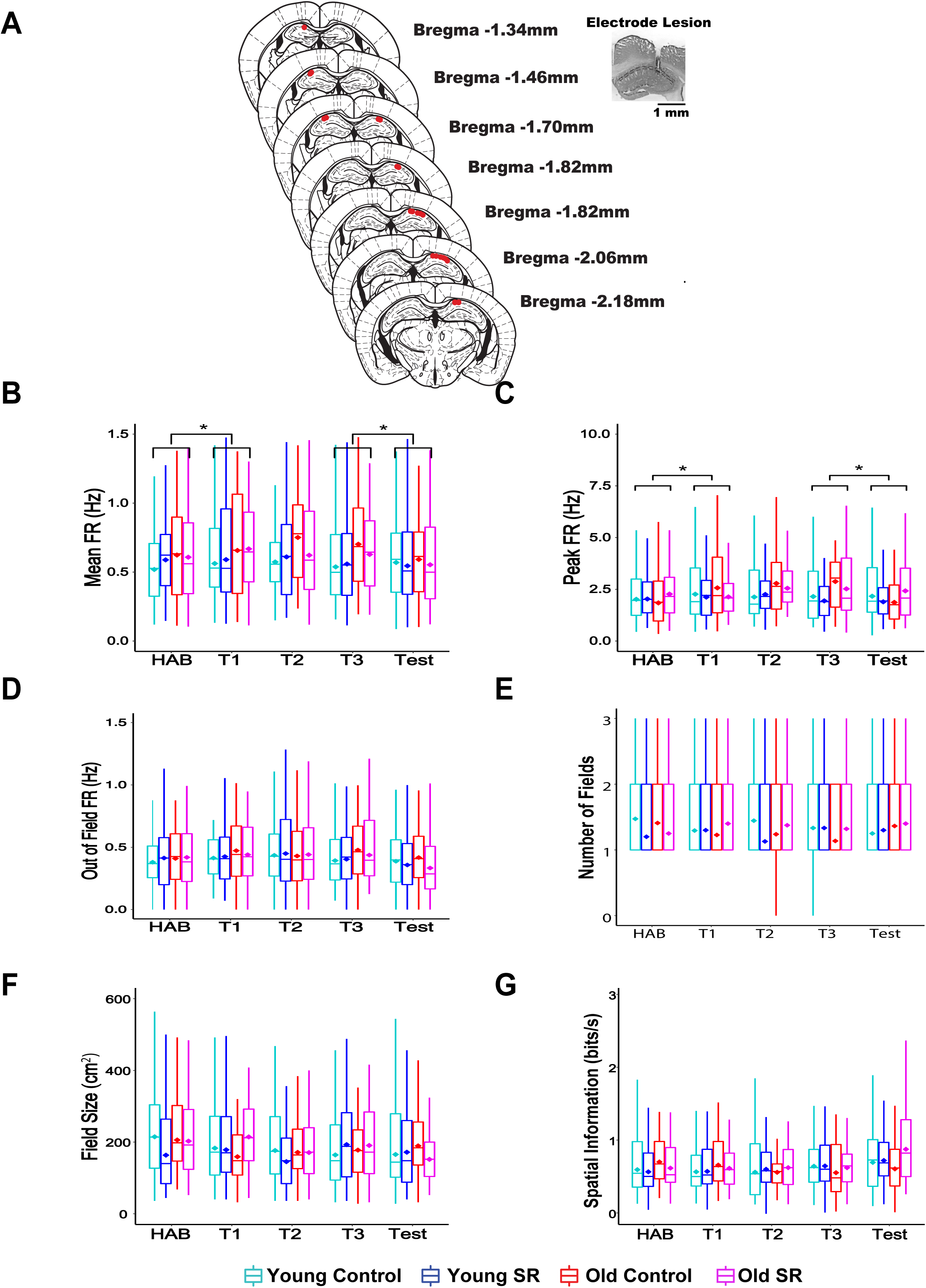
Schematic of electrode placements and place cell parameters recorded during OPR performance. A) Schematic of electrode placements (red dots) and microphotograph showing sample lesion marking electrode placement in CA1. B-D) Mean (B), peak (C), and out of field (D) firing rate for all groups across trials. There were no differences in mean or peak firing rate between the groups throughout training (p>0.05). However, there were increases in firing rate that persisted during training (T1 to T3) when the objects were introduced across all groups [MFR: Fw(4)=1.14, p<0.007; PFR: Fw(4)=1.11, p<0.02 . Analysis of simple effects indicated that all groups displayed higher mean and peak firing rates during the first object trial and test trials (MFR: Hab xT1: Zw=1.73, p<0.04; T3 x Test: Zw=3.16, p<0.04; PFR: Hab xT1: Zw=2.08, p<0.02; T3 x Test: Zw=0.96, p<0.05]. No differences were observed in out- of-field firing rate. E-G) There were no differences during training or testing in other place cell parameters, including number of fields (E), field size (F), and spatial information content (G, p>0.05). Hab: habituation, T1-T3: training trials. Asterisks (*) represent significance using alpha=0.05. Statistical Details in Supplemental Statistic-Tables.

**Supplemental Statistic-Tables.**
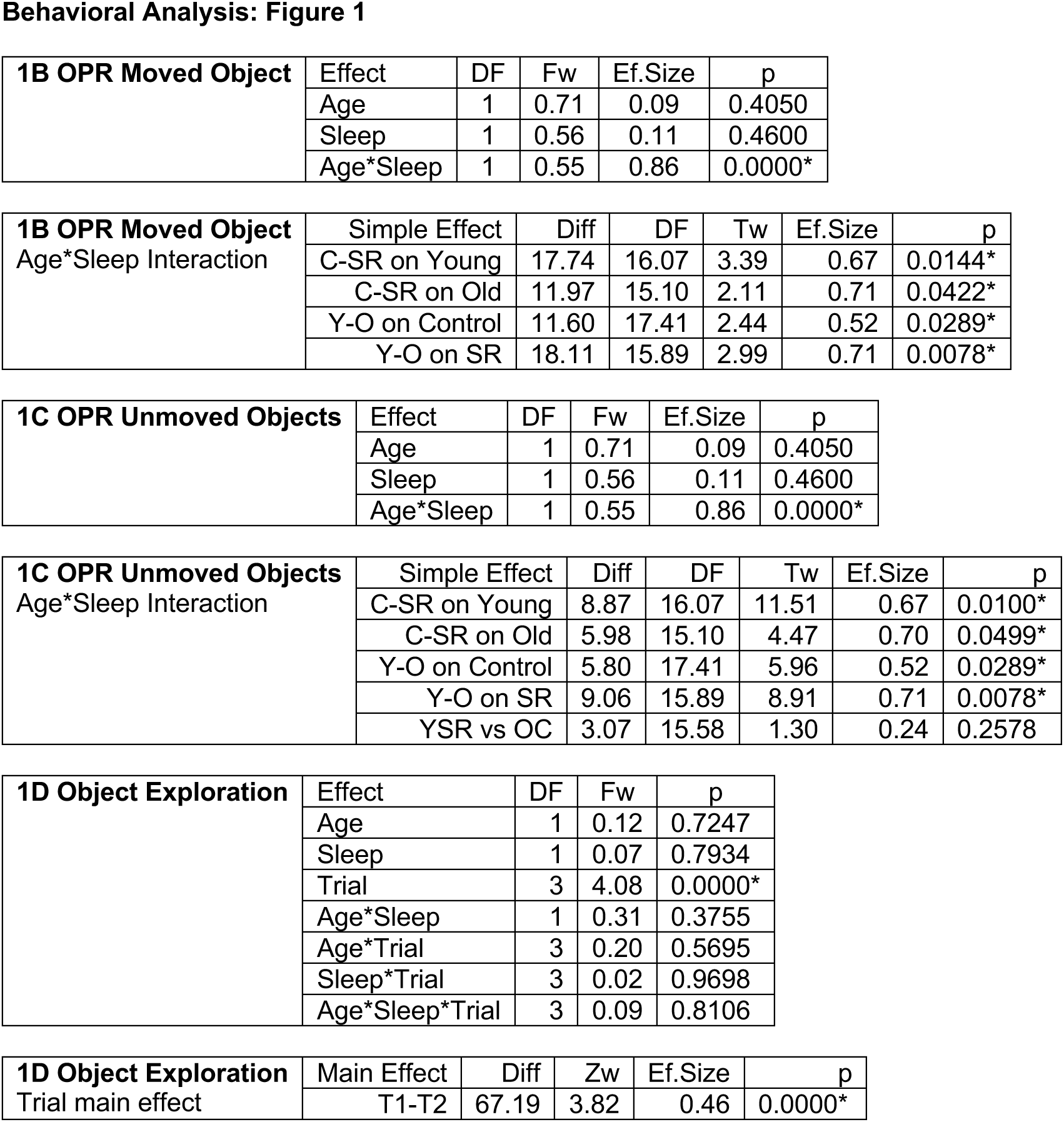
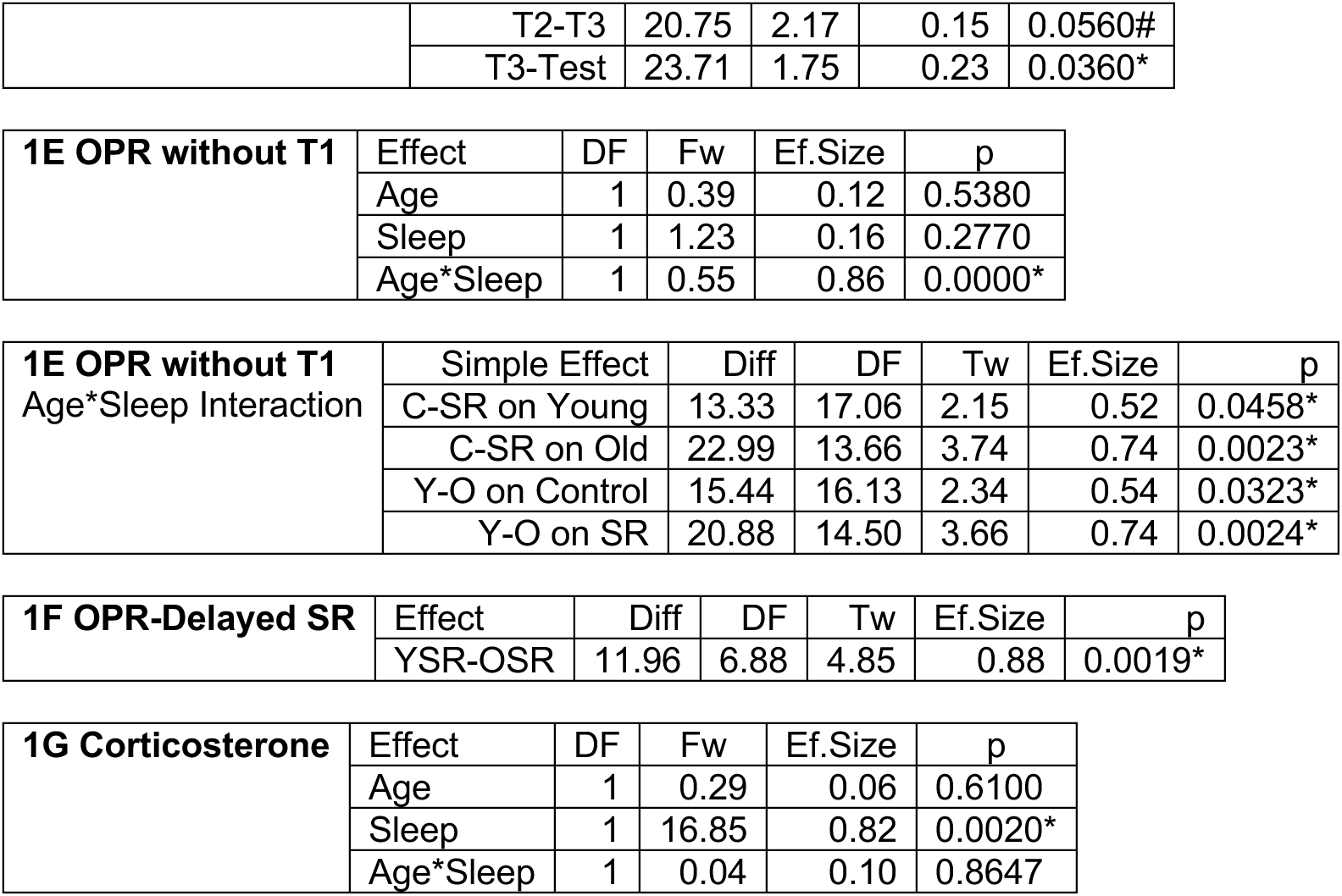
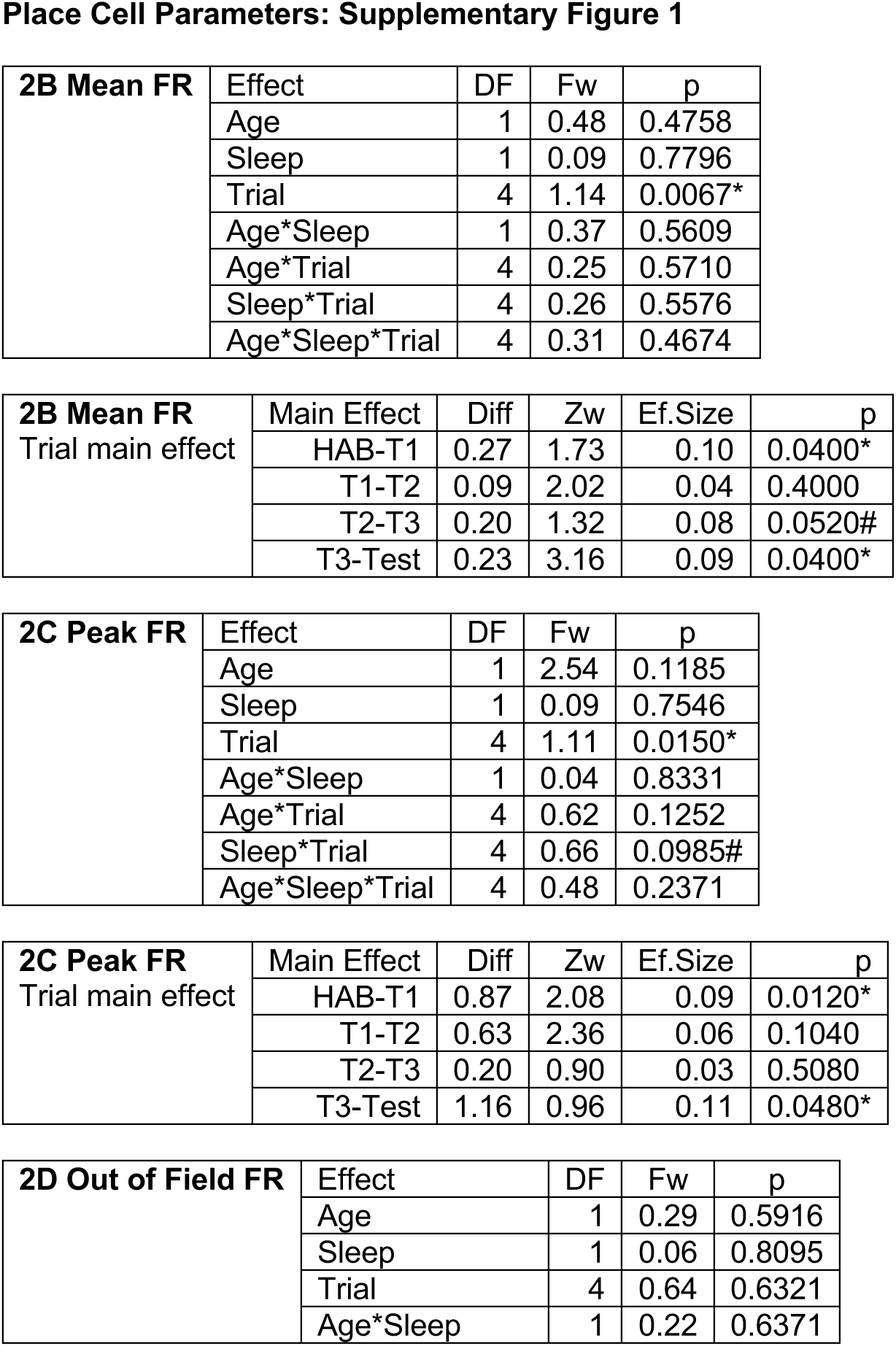
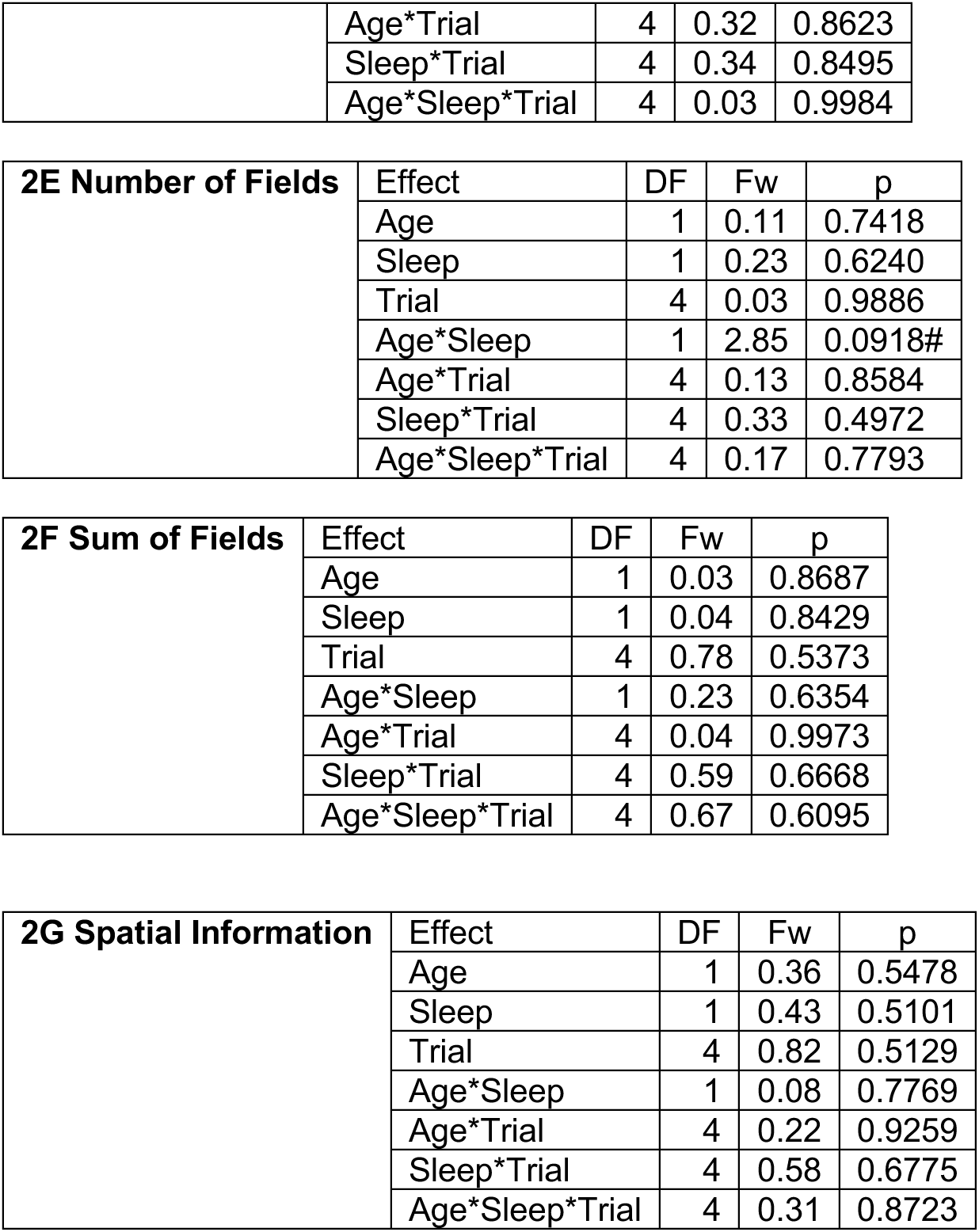
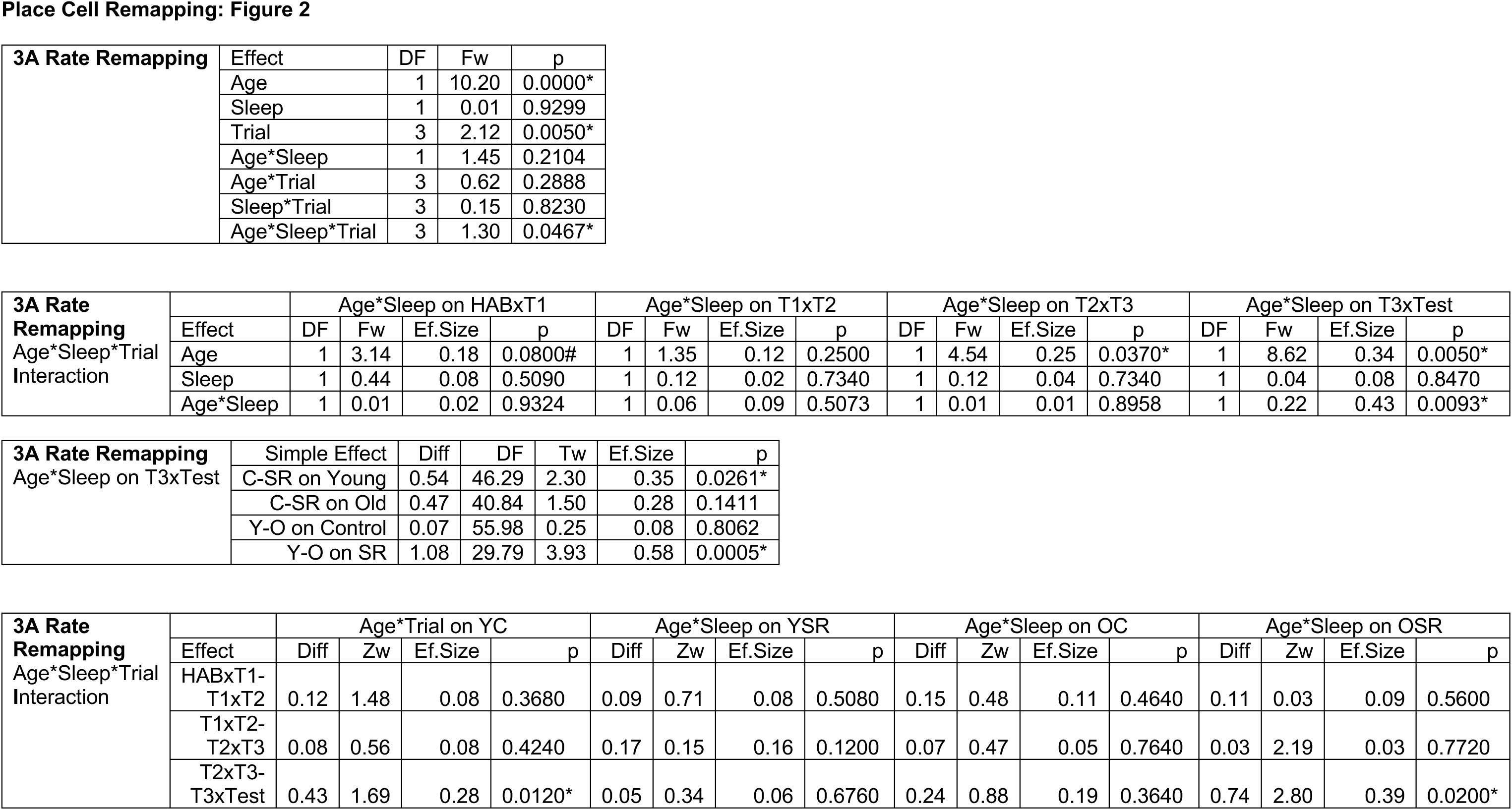
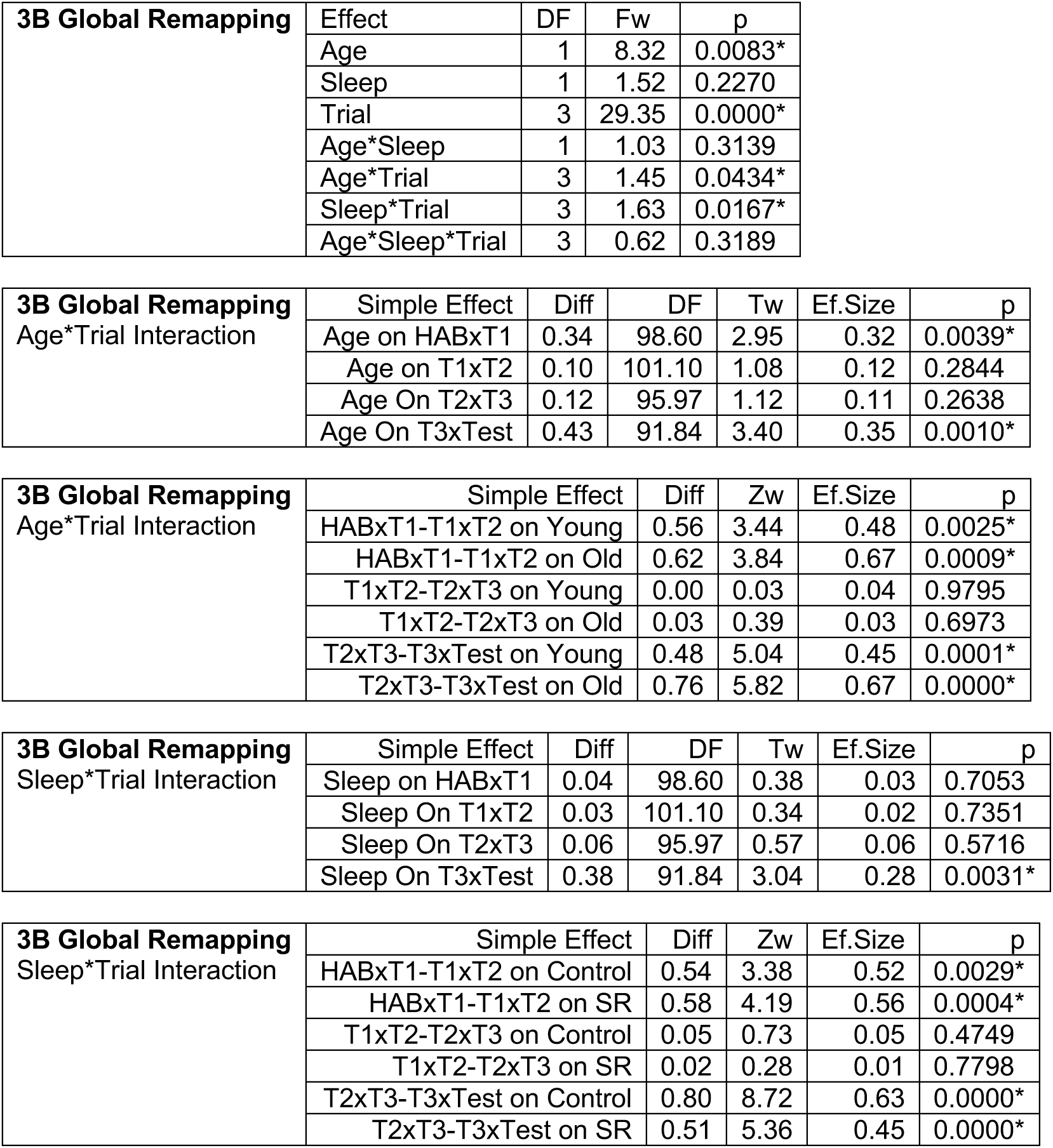
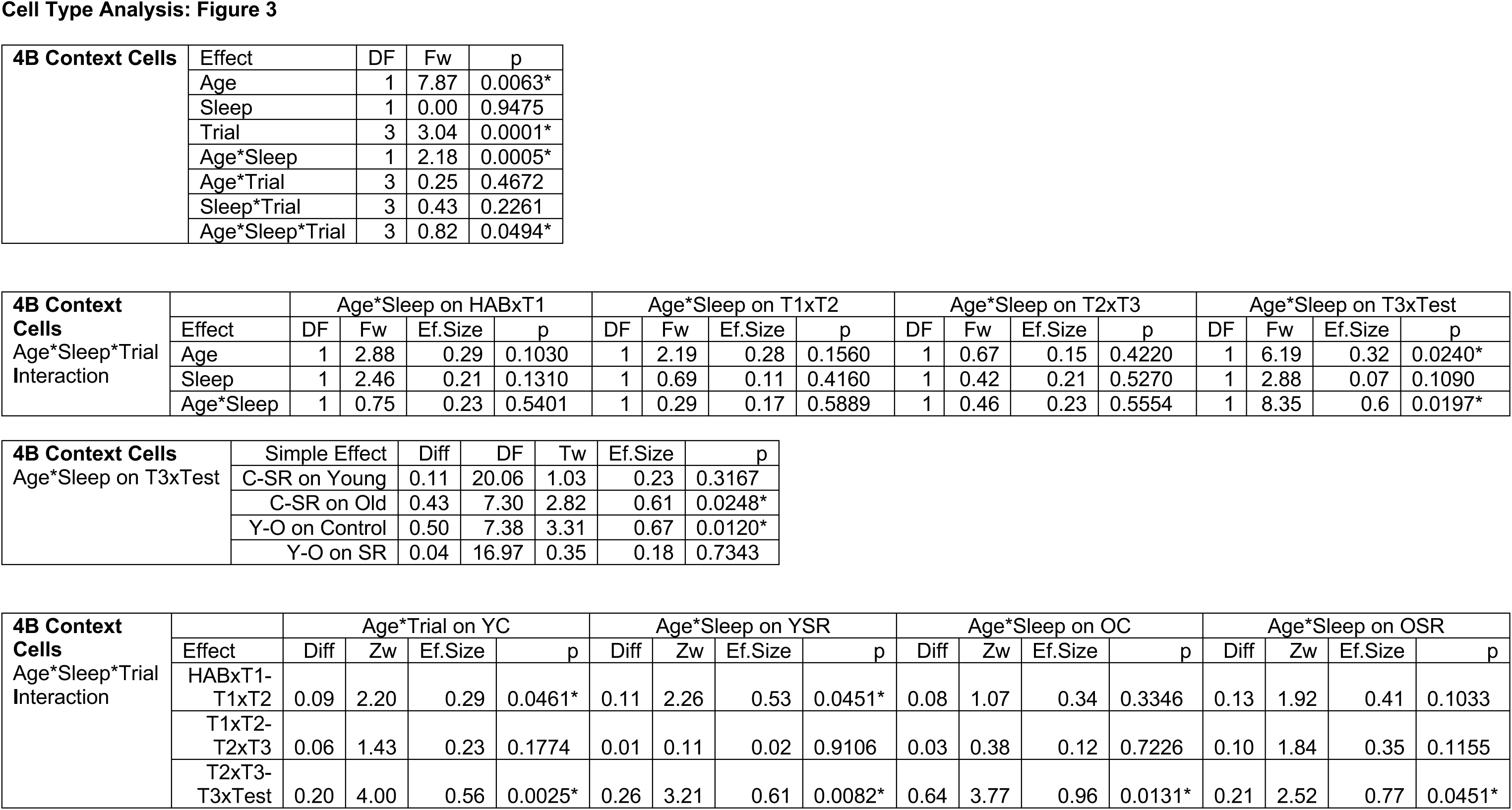
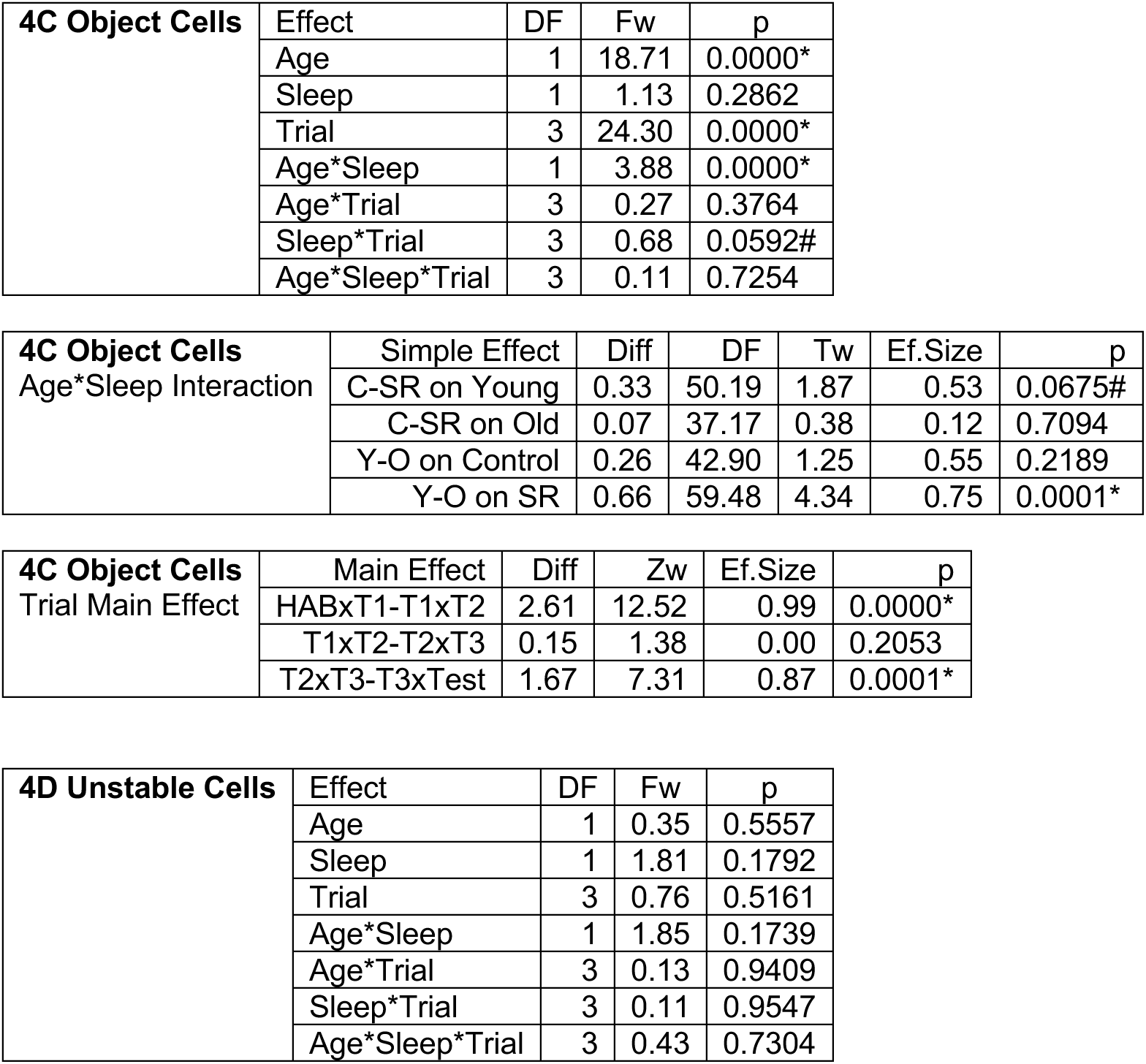
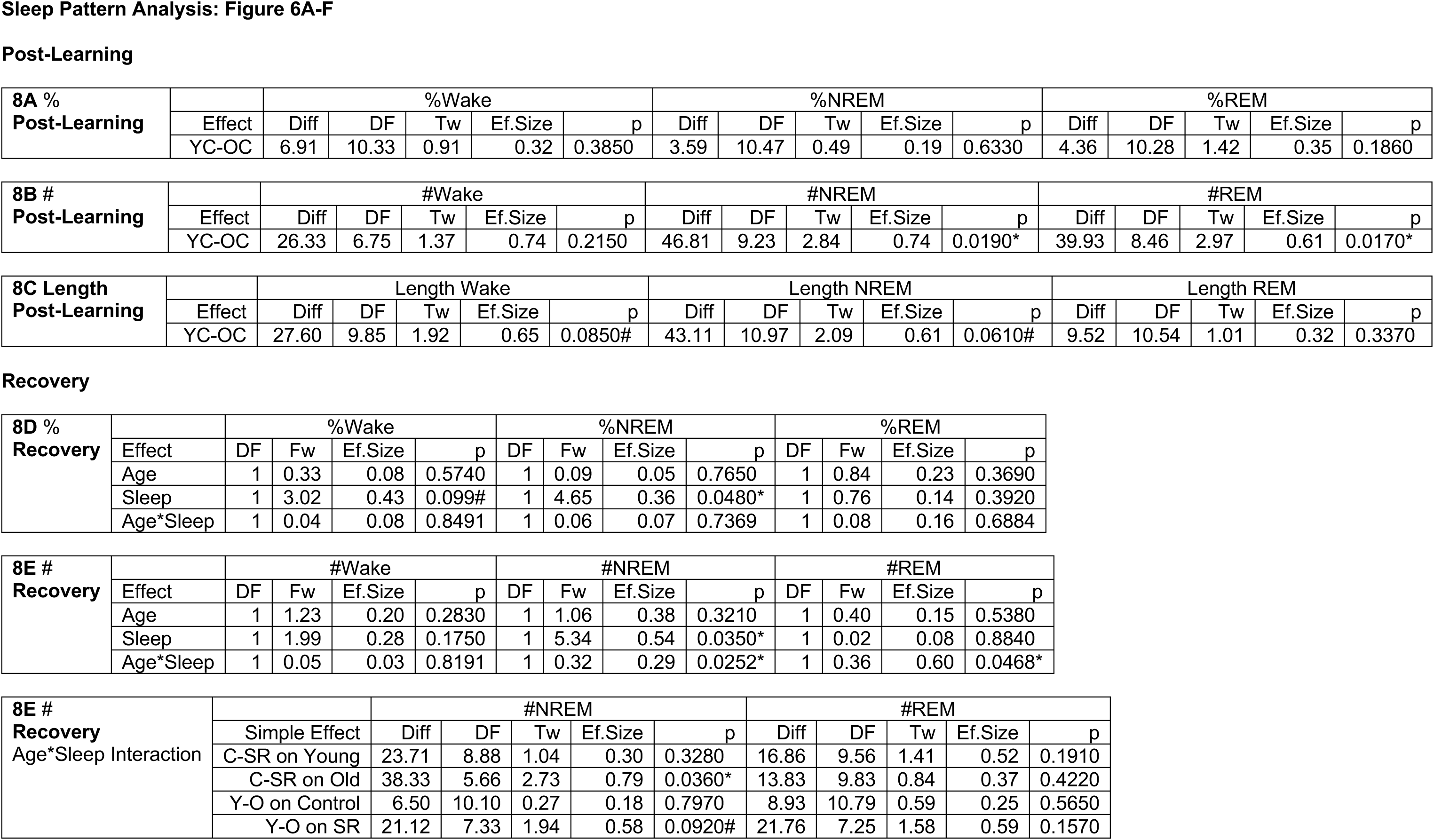
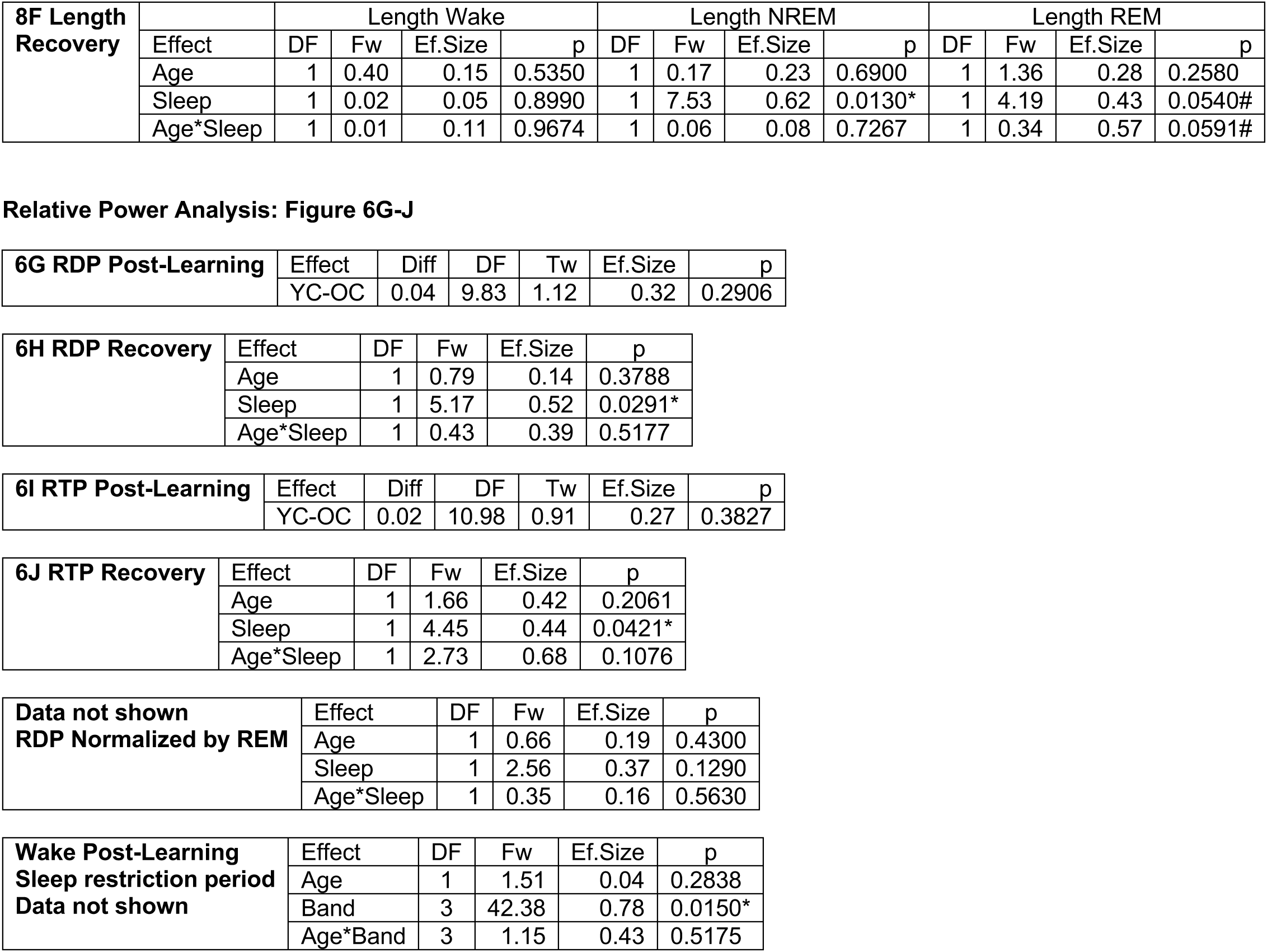
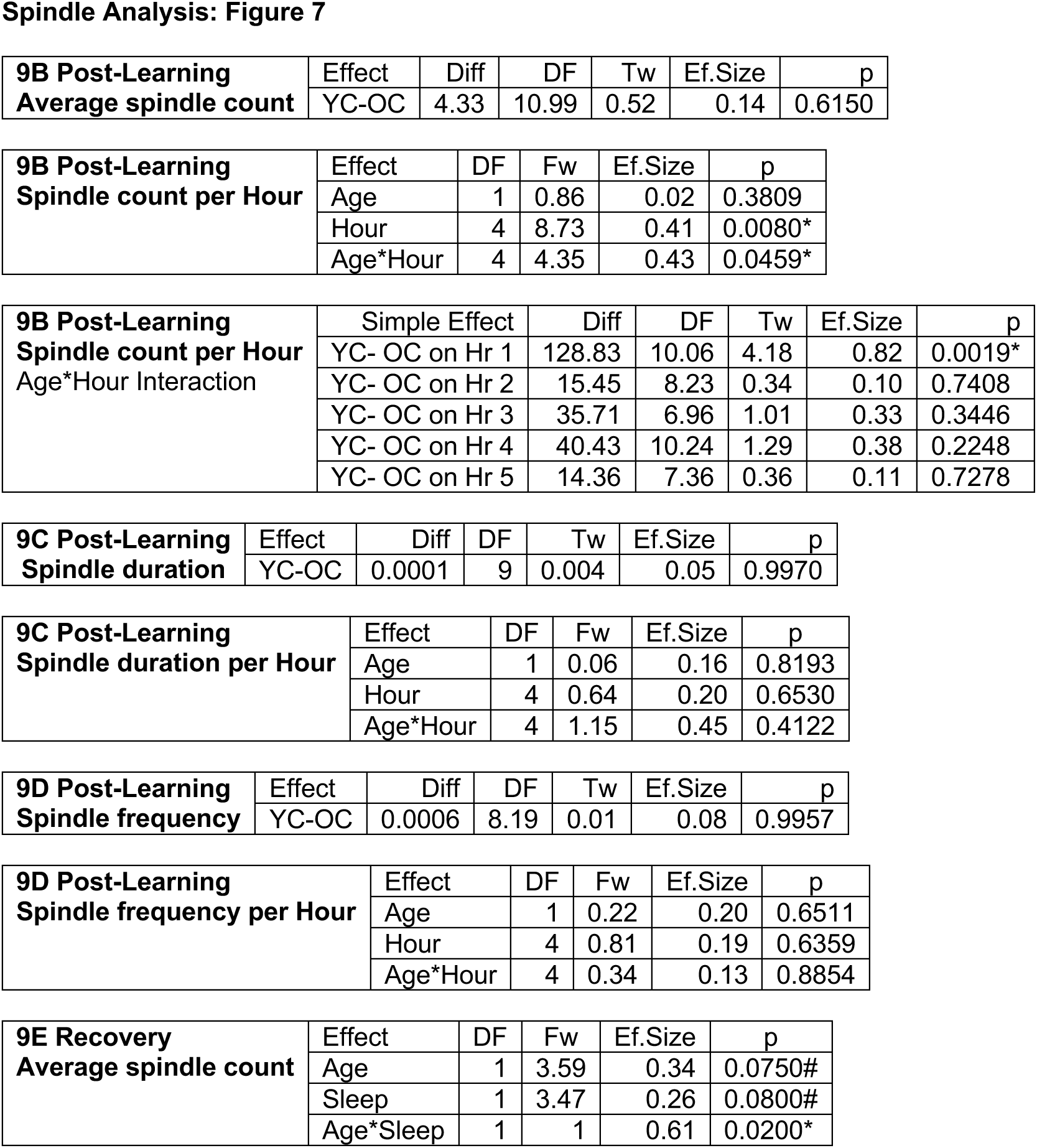
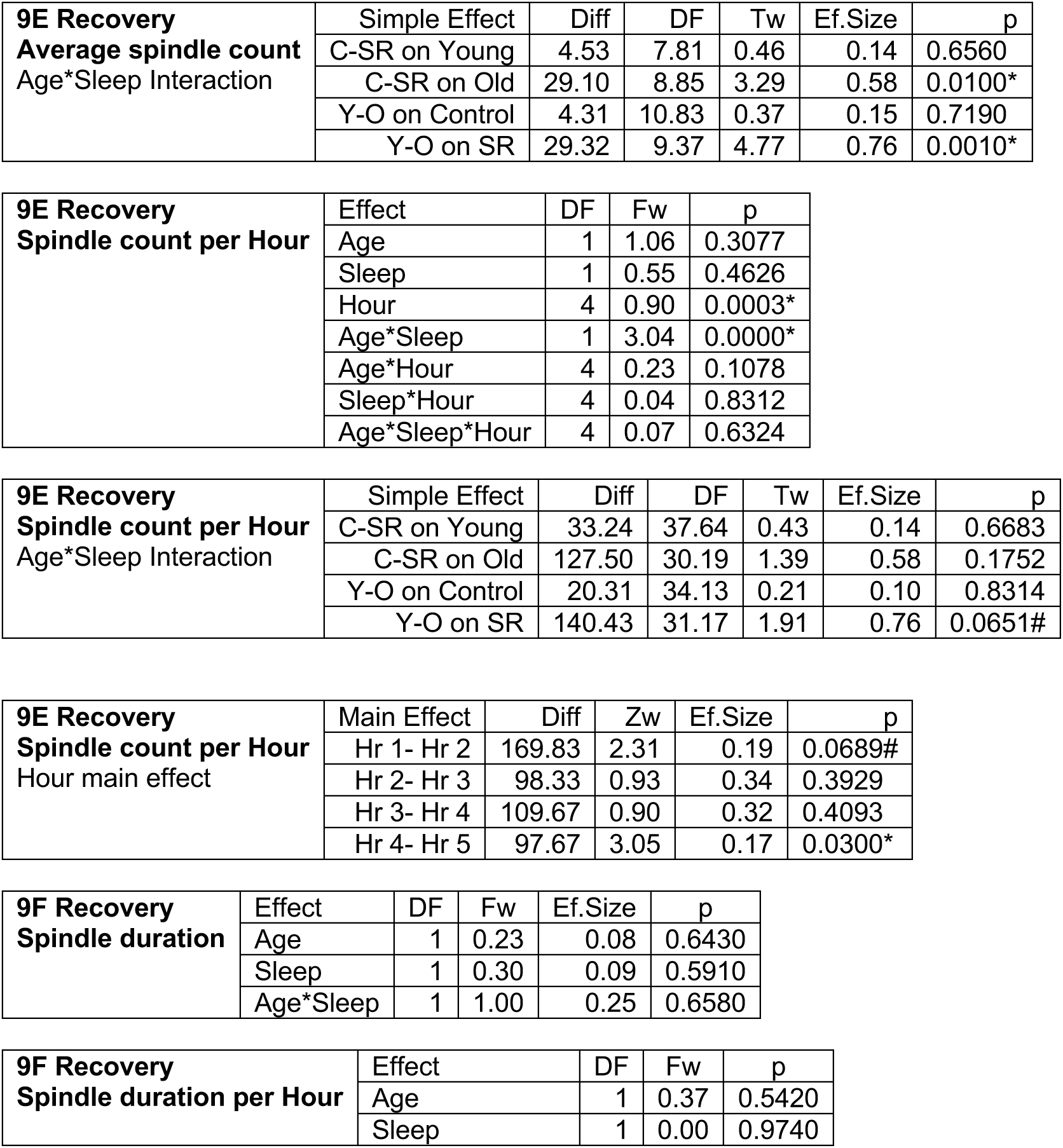
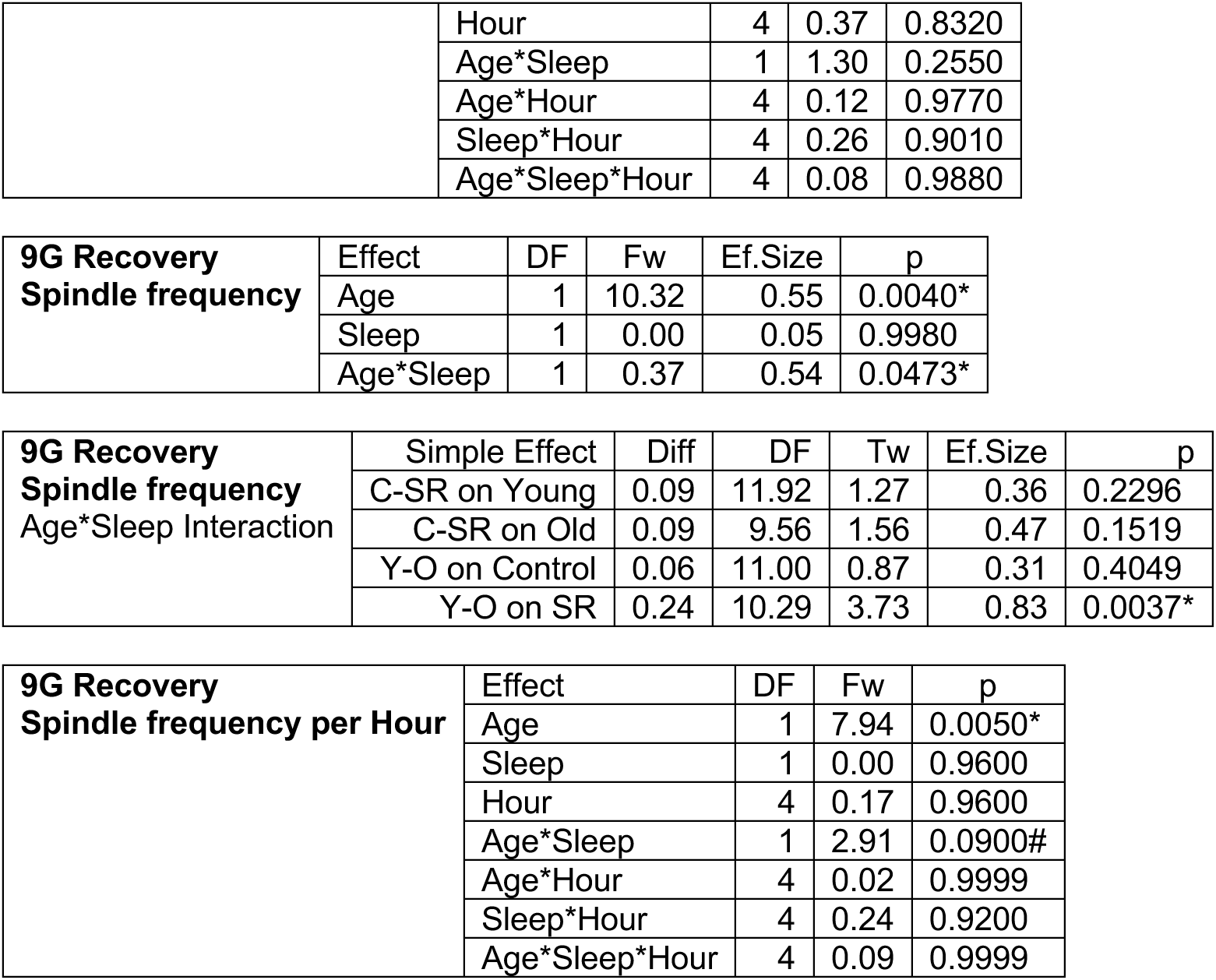

| **1B OPR Moved Object** | Effect | DF | Fw | Ef.Size | p |
| --- | --- | --- | --- | --- | --- |
|  | Age | 1 | 0.71 | 0.09 | 0.4050 |
|  | Sleep | 1 | 0.56 | 0.11 | 0.4600 |
|  | Age*Sleep | 1 | 0.55 | 0.86 | 0.0000* |

| **1B OPR Moved Object** | Simple Effect | Diff | DF | Tw | Ef.Size | p |
| --- | --- | --- | --- | --- | --- | --- |
| Age*Sleep Interaction | C-SR on Young | 17.74 | 16.07 | 3.39 | 0.67 | 0.0144* |
|  | C-SR on Old | 11.97 | 15.10 | 2.11 | 0.71 | 0.0422* |
|  | Y-O on Control | 11.60 | 17.41 | 2.44 | 0.52 | 0.0289* |
|  | Y-O on SR | 18.11 | 15.89 | 2.99 | 0.71 | 0.0078* |

| **1C OPR Unmoved Objects** | Effect | DF | Fw | Ef.Size | p |
| --- | --- | --- | --- | --- | --- |
|  | Age | 1 | 0.71 | 0.09 | 0.4050 |
|  | Sleep | 1 | 0.56 | 0.11 | 0.4600 |
|  | Age*Sleep | 1 | 0.55 | 0.86 | 0.0000* |

| **1C OPR Unmoved Objects** | Simple Effect | Diff | DF | Tw | Ef.Size | p |
| --- | --- | --- | --- | --- | --- | --- |
| Age*Sleep Interaction | C-SR on Young | 8.87 | 16.07 | 11.51 | 0.67 | 0.0100* |
|  | C-SR on Old | 5.98 | 15.10 | 4.47 | 0.70 | 0.0499* |
|  | Y-O on Control | 5.80 | 17.41 | 5.96 | 0.52 | 0.0289* |
|  | Y-O on SR | 9.06 | 15.89 | 8.91 | 0.71 | 0.0078* |
|  | YSR vs OC | 3.07 | 15.58 | 1.30 | 0.24 | 0.2578 |

| **1D Object Exploration** | Effect | DF | Fw | p |
| --- | --- | --- | --- | --- |
|  | Age | 1 | 0.12 | 0.7247 |
|  | Sleep | 1 | 0.07 | 0.7934 |
|  | Trial | 3 | 4.08 | 0.0000* |
|  | Age*Sleep | 1 | 0.31 | 0.3755 |
|  | Age*Trial | 3 | 0.20 | 0.5695 |
|  | Sleep*Trial | 3 | 0.02 | 0.9698 |
|  | Age*Sleep*Trial | 3 | 0.09 | 0.8106 |

| **1D Object Exploration** | Main Effect | Diff | Zw | Ef.Size | p |
| --- | --- | --- | --- | --- | --- |
| Trial main effect | T1-T2 | 67.19 | 3.82 | 0.46 | 0.0000* |
|  | T2-T3 | 20.75 | 2.17 | 0.15 | 0.0560# |
|  | T3-Test | 23.71 | 1.75 | 0.23 | 0.0360* |

| 1E OPR without T1 | Effect | DF | Fw | Ef.Size | p |
| --- | --- | --- | --- | --- | --- |
|  | Age | 1 | 0.39 | 0.12 | 0.5380 |
|  | Sleep | 1 | 1.23 | 0.16 | 0.2770 |
|  | Age*Sleep | 1 | 0.55 | 0.86 | 0.0000* |

| 1E OPR without T1 | Simple Effect | Diff | DF | Tw | Ef.Size | p |
| --- | --- | --- | --- | --- | --- | --- |
| Age*Sleep Interaction | C-SR on Young | 13.33 | 17.06 | 2.15 | 0.52 | 0.0458* |
|  | C-SR on Old | 22.99 | 13.66 | 3.74 | 0.74 | 0.0023* |
|  | Y-O on Control | 15.44 | 16.13 | 2.34 | 0.54 | 0.0323* |
|  | Y-O on SR | 20.88 | 14.50 | 3.66 | 0.74 | 0.0024* |

| 1F OPR-Delayed SR | Effect | Diff | DF | Tw | Ef.Size | p |
| --- | --- | --- | --- | --- | --- | --- |
|  | YSR-OSR | 11.96 | 6.88 | 4.85 | 0.88 | 0.0019* |

| 1G Corticosterone | Effect | DF | Fw | Ef.Size | p |
| --- | --- | --- | --- | --- | --- |
|  | Age | 1 | 0.29 | 0.06 | 0.6100 |
|  | Sleep | 1 | 16.85 | 0.82 | 0.0020* |
|  | Age*Sleep | 1 | 0.04 | 0.10 | 0.8647 |

Place Cell Parameters: Supplementary Figure 1
| 2B Mean FR | Effect | DF | Fw | p |
| --- | --- | --- | --- | --- |
|  | Age | 1 | 0.48 | 0.4758 |
|  | Sleep | 1 | 0.09 | 0.7796 |
|  | Trial | 4 | 1.14 | 0.0067* |
|  | Age*Sleep | 1 | 0.37 | 0.5609 |
|  | Age*Trial | 4 | 0.25 | 0.5710 |
|  | Sleep*Trial | 4 | 0.26 | 0.5576 |
|  | Age*Sleep*Trial | 4 | 0.31 | 0.4674 |

| 2B Mean FR | Main Effect | Diff | Zw | Ef.Size | p |
| --- | --- | --- | --- | --- | --- |
| Trial main effect | HAB-T1 | 0.27 | 1.73 | 0.10 | 0.0400* |
|  | T1-T2 | 0.09 | 2.02 | 0.04 | 0.4000 |
|  | T2-T3 | 0.20 | 1.32 | 0.08 | 0.0520# |
|  | T3-Test | 0.23 | 3.16 | 0.09 | 0.0400* |

| 2C Peak FR | Effect | DF | Fw | p |
| --- | --- | --- | --- | --- |
|  | Age | 1 | 2.54 | 0.1185 |
|  | Sleep | 1 | 0.09 | 0.7546 |
|  | Trial | 4 | 1.11 | 0.0150* |
|  | Age*Sleep | 1 | 0.04 | 0.8331 |
|  | Age*Trial | 4 | 0.62 | 0.1252 |
|  | Sleep*Trial | 4 | 0.66 | 0.0985# |
|  | Age*Sleep*Trial | 4 | 0.48 | 0.2371 |

| 2C Peak FR | Main Effect | Diff | Zw | Ef.Size | p |
| --- | --- | --- | --- | --- | --- |
| Trial main effect | HAB-T1 | 0.87 | 2.08 | 0.09 | 0.0120* |
|  | T1-T2 | 0.63 | 2.36 | 0.06 | 0.1040 |
|  | T2-T3 | 0.20 | 0.90 | 0.03 | 0.5080 |
|  | T3-Test | 1.16 | 0.96 | 0.11 | 0.0480* |

| 2D Out of Field FR | Effect | DF | Fw | p |
| --- | --- | --- | --- | --- |
|  | Age | 1 | 0.29 | 0.5916 |
|  | Sleep | 1 | 0.06 | 0.8095 |
|  | Trial | 4 | 0.64 | 0.6321 |
|  | Age*Sleep | 1 | 0.22 | 0.6371 |
|  | Age*Trial | 4 | 0.32 | 0.8623 |
|  | Sleep*Trial | 4 | 0.34 | 0.8495 |
|  | Age*Sleep*Trial | 4 | 0.03 | 0.9984 |

| 2E Number of Fields | Effect | DF | Fw | p |
| --- | --- | --- | --- | --- |
|  | Age | 1 | 0.11 | 0.7418 |
|  | Sleep | 1 | 0.23 | 0.6240 |
|  | Trial | 4 | 0.03 | 0.9886 |
|  | Age*Sleep | 1 | 2.85 | 0.0918# |
|  | Age*Trial | 4 | 0.13 | 0.8584 |
|  | Sleep*Trial | 4 | 0.33 | 0.4972 |
|  | Age*Sleep*Trial | 4 | 0.17 | 0.7793 |

| 2F Sum of Fields | Effect | DF | Fw | p |
| --- | --- | --- | --- | --- |
|  | Age | 1 | 0.03 | 0.8687 |
|  | Sleep | 1 | 0.04 | 0.8429 |
|  | Trial | 4 | 0.78 | 0.5373 |
|  | Age*Sleep | 1 | 0.23 | 0.6354 |
|  | Age*Trial | 4 | 0.04 | 0.9973 |
|  | Sleep*Trial | 4 | 0.59 | 0.6668 |
|  | Age*Sleep*Trial | 4 | 0.67 | 0.6095 |

| 2G Spatial Information | Effect | DF | Fw | p |
| --- | --- | --- | --- | --- |
|  | Age | 1 | 0.36 | 0.5478 |
|  | Sleep | 1 | 0.43 | 0.5101 |
|  | Trial | 4 | 0.82 | 0.5129 |
|  | Age*Sleep | 1 | 0.08 | 0.7769 |
|  | Age*Trial | 4 | 0.22 | 0.9259 |
|  | Sleep*Trial | 4 | 0.58 | 0.6775 |
|  | Age*Sleep*Trial | 4 | 0.31 | 0.8723 |

Place Cell Remapping: Figure 2
| 3A Rate Remapping | Effect | DF | Fw | p |
| --- | --- | --- | --- | --- |
|  | Age | 1 | 10.20 | 0.0000* |
|  | Sleep | 1 | 0.01 | 0.9299 |
|  | Trial | 3 | 2.12 | 0.0050* |
|  | Age*Sleep | 1 | 1.45 | 0.2104 |
|  | Age*Trial | 3 | 0.62 | 0.2888 |
|  | Sleep*Trial | 3 | 0.15 | 0.8230 |
|  | Age*Sleep*Trial | 3 | 1.30 | 0.0467* |

| 3A Rate Remapping<br>Age*Sleep*Trial<br>Interaction |  | Age*Sleep on HABxT1 |  |  |  | Age*Sleep on T1xT2 |  |  |  | Age*Sleep on T2xT3 |  |  |  | Age*Sleep on T3xTest |  |  |  |
| --- | --- | --- | --- | --- | --- | --- | --- | --- | --- | --- | --- | --- | --- | --- | --- | --- | --- |
|  | Effect | DF | Fw | Ef.Size | p | DF | Fw | Ef.Size | p | DF | Fw | Ef.Size | p | DF | Fw | Ef.Size | p |
|  | Age | 1 | 3.14 | 0.18 | 0.0800# | 1 | 1.35 | 0.12 | 0.2500 | 1 | 4.54 | 0.25 | 0.0370* | 1 | 8.62 | 0.34 | 0.0050* |
|  | Sleep | 1 | 0.44 | 0.08 | 0.5090 | 1 | 0.12 | 0.02 | 0.7340 | 1 | 0.12 | 0.04 | 0.7340 | 1 | 0.04 | 0.08 | 0.8470 |
|  | Age*Sleep | 1 | 0.01 | 0.02 | 0.9324 | 1 | 0.06 | 0.09 | 0.5073 | 1 | 0.01 | 0.01 | 0.8958 | 1 | 0.22 | 0.43 | 0.0093* |

| 3A Rate Remapping<br>Age*Sleep on T3xTest | Simple Effect | Diff | DF | Tw | Ef.Size | p |
| --- | --- | --- | --- | --- | --- | --- |
|  | C-SR on Young | 0.54 | 46.29 | 2.30 | 0.35 | 0.0261* |
|  | C-SR on Old | 0.47 | 40.84 | 1.50 | 0.28 | 0.1411 |
|  | Y-O on Control | 0.07 | 55.98 | 0.25 | 0.08 | 0.8062 |
|  | Y-O on SR | 1.08 | 29.79 | 3.93 | 0.58 | 0.0005* |

| 3A Rate Remapping<br>Age*Sleep*Trial<br>Interaction |  | Age*Trial on YC |  |  |  | Age*Sleep on YSR |  |  |  | Age*Sleep on OC |  |  |  | Age*Sleep on OSR |  |  |  |
| --- | --- | --- | --- | --- | --- | --- | --- | --- | --- | --- | --- | --- | --- | --- | --- | --- | --- |
|  | Effect | Diff | Zw | Ef.Size | p | Diff | Zw | Ef.Size | p | Diff | Zw | Ef.Size | p | Diff | Zw | Ef.Size | p |
|  | HABxT1-T1xT2 | 0.12 | 1.48 | 0.08 | 0.3680 | 0.09 | 0.71 | 0.08 | 0.5080 | 0.15 | 0.48 | 0.11 | 0.4640 | 0.11 | 0.03 | 0.09 | 0.5600 |
|  | T1xT2-T2xT3 | 0.08 | 0.56 | 0.08 | 0.4240 | 0.17 | 0.15 | 0.16 | 0.1200 | 0.07 | 0.47 | 0.05 | 0.7640 | 0.03 | 2.19 | 0.03 | 0.7720 |
|  | T2xT3-T3xTest | 0.43 | 1.69 | 0.28 | 0.0120* | 0.05 | 0.34 | 0.06 | 0.6760 | 0.24 | 0.88 | 0.19 | 0.3640 | 0.74 | 2.80 | 0.39 | 0.0200* |

| 3B Global Remapping | Effect | DF | Fw | p |
| --- | --- | --- | --- | --- |
|  | Age | 1 | 8.32 | 0.0083* |
|  | Sleep | 1 | 1.52 | 0.2270 |
|  | Trial | 3 | 29.35 | 0.0000* |
|  | Age*Sleep | 1 | 1.03 | 0.3139 |
|  | Age*Trial | 3 | 1.45 | 0.0434* |
|  | Sleep*Trial | 3 | 1.63 | 0.0167* |
|  | Age*Sleep*Trial | 3 | 0.62 | 0.3189 |

| 3B Global Remapping<br>Age*Trial Interaction | Simple Effect | Diff | DF | Tw | Ef.Size | p |
| --- | --- | --- | --- | --- | --- | --- |
|  | Age on HABxT1 | 0.34 | 98.60 | 2.95 | 0.32 | 0.0039* |
|  | Age on T1xT2 | 0.10 | 101.10 | 1.08 | 0.12 | 0.2844 |
|  | Age On T2xT3 | 0.12 | 95.97 | 1.12 | 0.11 | 0.2638 |
|  | Age On T3xTest | 0.43 | 91.84 | 3.40 | 0.35 | 0.0010* |

| 3B Global Remapping<br>Age*Trial Interaction | Simple Effect | Diff | Zw | Ef.Size | p |
| --- | --- | --- | --- | --- | --- |
|  | HABxT1-T1xT2 on Young | 0.56 | 3.44 | 0.48 | 0.0025* |
|  | HABxT1-T1xT2 on Old | 0.62 | 3.84 | 0.67 | 0.0009* |
|  | T1xT2-T2xT3 on Young | 0.00 | 0.03 | 0.04 | 0.9795 |
|  | T1xT2-T2xT3 on Old | 0.03 | 0.39 | 0.03 | 0.6973 |
|  | T2xT3-T3xTest on Young | 0.48 | 5.04 | 0.45 | 0.0001* |
|  | T2xT3-T3xTest on Old | 0.76 | 5.82 | 0.67 | 0.0000* |

| 3B Global Remapping<br>Sleep*Trial Interaction | Simple Effect | Diff | DF | Tw | Ef.Size | p |
| --- | --- | --- | --- | --- | --- | --- |
|  | Sleep on HABxT1 | 0.04 | 98.60 | 0.38 | 0.03 | 0.7053 |
|  | Sleep On T1xT2 | 0.03 | 101.10 | 0.34 | 0.02 | 0.7351 |
|  | Sleep On T2xT3 | 0.06 | 95.97 | 0.57 | 0.06 | 0.5716 |
|  | Sleep On T3xTest | 0.38 | 91.84 | 3.04 | 0.28 | 0.0031* |

| 3B Global Remapping<br>Sleep*Trial Interaction | Simple Effect | Diff | Zw | Ef.Size | p |
| --- | --- | --- | --- | --- | --- |
|  | HABxT1-T1xT2 on Control | 0.54 | 3.38 | 0.52 | 0.0029* |
|  | HABxT1-T1xT2 on SR | 0.58 | 4.19 | 0.56 | 0.0004* |
|  | T1xT2-T2xT3 on Control | 0.05 | 0.73 | 0.05 | 0.4749 |
|  | T1xT2-T2xT3 on SR | 0.02 | 0.28 | 0.01 | 0.7798 |
|  | T2xT3-T3xTest on Control | 0.80 | 8.72 | 0.63 | 0.0000* |
|  | T2xT3-T3xTest on SR | 0.51 | 5.36 | 0.45 | 0.0000* |

Cell Type Analysis: Figure 3
| 4B Context Cells | Effect | DF | Fw | p |
| --- | --- | --- | --- | --- |
|  | Age | 1 | 7.87 | 0.0063* |
|  | Sleep | 1 | 0.00 | 0.9475 |
|  | Trial | 3 | 3.04 | 0.0001* |
|  | Age*Sleep | 1 | 2.18 | 0.0005* |
|  | Age*Trial | 3 | 0.25 | 0.4672 |
|  | Sleep*Trial | 3 | 0.43 | 0.2261 |
|  | Age*Sleep*Trial | 3 | 0.82 | 0.0494* |

| 4B Context Cells<br>Age*Sleep*Trial Interaction |  | Age*Sleep on HABxT1 |  |  |  | Age*Sleep on T1xT2 |  |  |  | Age*Sleep on T2xT3 |  |  |  | Age*Sleep on T3xTest |  |  |  |
| --- | --- | --- | --- | --- | --- | --- | --- | --- | --- | --- | --- | --- | --- | --- | --- | --- | --- |
|  | Effect | DF | Fw | Ef.Size | p | DF | Fw | Ef.Size | p | DF | Fw | Ef.Size | p | DF | Fw | Ef.Size | p |
|  | Age | 1 | 2.88 | 0.29 | 0.1030 | 1 | 2.19 | 0.28 | 0.1560 | 1 | 0.67 | 0.15 | 0.4220 | 1 | 6.19 | 0.32 | 0.0240* |
|  | Sleep | 1 | 2.46 | 0.21 | 0.1310 | 1 | 0.69 | 0.11 | 0.4160 | 1 | 0.42 | 0.21 | 0.5270 | 1 | 2.88 | 0.07 | 0.1090 |
|  | Age*Sleep | 1 | 0.75 | 0.23 | 0.5401 | 1 | 0.29 | 0.17 | 0.5889 | 1 | 0.46 | 0.23 | 0.5554 | 1 | 8.35 | 0.6 | 0.0197* |

| 4B Context Cells<br>Age*Sleep on T3xTest | Simple Effect | Diff | DF | Tw | Ef.Size | p |
| --- | --- | --- | --- | --- | --- | --- |
|  | C-SR on Young | 0.11 | 20.06 | 1.03 | 0.23 | 0.3167 |
|  | C-SR on Old | 0.43 | 7.30 | 2.82 | 0.61 | 0.0248* |
|  | Y-O on Control | 0.50 | 7.38 | 3.31 | 0.67 | 0.0120* |
|  | Y-O on SR | 0.04 | 16.97 | 0.35 | 0.18 | 0.7343 |

| 4B Context Cells<br>Age*Sleep*Trial Interaction |  | Age*Trial on YC |  |  |  | Age*Sleep on YSR |  |  |  | Age*Sleep on OC |  |  |  | Age*Sleep on OSR |  |  |  |
| --- | --- | --- | --- | --- | --- | --- | --- | --- | --- | --- | --- | --- | --- | --- | --- | --- | --- |
|  | Effect | Diff | Zw | Ef.Size | p | Diff | Zw | Ef.Size | p | Diff | Zw | Ef.Size | p | Diff | Zw | Ef.Size | p |
|  | HABxT1-T1xT2 | 0.09 | 2.20 | 0.29 | 0.0461* | 0.11 | 2.26 | 0.53 | 0.0451* | 0.08 | 1.07 | 0.34 | 0.3346 | 0.13 | 1.92 | 0.41 | 0.1033 |
|  | T1xT2-T2xT3 | 0.06 | 1.43 | 0.23 | 0.1774 | 0.01 | 0.11 | 0.02 | 0.9106 | 0.03 | 0.38 | 0.12 | 0.7226 | 0.10 | 1.84 | 0.35 | 0.1155 |
|  | T2xT3-T3xTest | 0.20 | 4.00 | 0.56 | 0.0025* | 0.26 | 3.21 | 0.61 | 0.0082* | 0.64 | 3.77 | 0.96 | 0.0131* | 0.21 | 2.52 | 0.77 | 0.0451* |

| 4C Object Cells | Effect | DF | Fw | p |
| --- | --- | --- | --- | --- |
|  | Age | 1 | 18.71 | 0.0000* |
|  | Sleep | 1 | 1.13 | 0.2862 |
|  | Trial | 3 | 24.30 | 0.0000* |
|  | Age*Sleep | 1 | 3.88 | 0.0000* |
|  | Age*Trial | 3 | 0.27 | 0.3764 |
|  | Sleep*Trial | 3 | 0.68 | 0.0592# |
|  | Age*Sleep*Trial | 3 | 0.11 | 0.7254 |

| 4C Object Cells | Simple Effect | Diff | DF | Tw | Ef.Size | p |
| --- | --- | --- | --- | --- | --- | --- |
| Age*Sleep Interaction | C-SR on Young | 0.33 | 50.19 | 1.87 | 0.53 | 0.0675# |
|  | C-SR on Old | 0.07 | 37.17 | 0.38 | 0.12 | 0.7094 |
|  | Y-O on Control | 0.26 | 42.90 | 1.25 | 0.55 | 0.2189 |
|  | Y-O on SR | 0.66 | 59.48 | 4.34 | 0.75 | 0.0001* |

| 4C Object Cells | Main Effect | Diff | Zw | Ef.Size | p |
| --- | --- | --- | --- | --- | --- |
| Trial Main Effect | HABxT1-T1xT2 | 2.61 | 12.52 | 0.99 | 0.0000* |
|  | T1xT2-T2xT3 | 0.15 | 1.38 | 0.00 | 0.2053 |
|  | T2xT3-T3xTest | 1.67 | 7.31 | 0.87 | 0.0001* |

| 4D Unstable Cells | Effect | DF | Fw | p |
| --- | --- | --- | --- | --- |
|  | Age | 1 | 0.35 | 0.5557 |
|  | Sleep | 1 | 1.81 | 0.1792 |
|  | Trial | 3 | 0.76 | 0.5161 |
|  | Age*Sleep | 1 | 1.85 | 0.1739 |
|  | Age*Trial | 3 | 0.13 | 0.9409 |
|  | Sleep*Trial | 3 | 0.11 | 0.9547 |
|  | Age*Sleep*Trial | 3 | 0.43 | 0.7304 |

Sleep Pattern Analysis: Figure 6A-F Post-Learning
| 8A %<br>Post-Learning |  | %Wake |  |  |  |  | %NREM |  |  |  |  | %REM |  |  |  |  |
| --- | --- | --- | --- | --- | --- | --- | --- | --- | --- | --- | --- | --- | --- | --- | --- | --- |
|  | Effect | Diff | DF | Tw | Ef.Size | p | Diff | DF | Tw | Ef.Size | p | Diff | DF | Tw | Ef.Size | p |
|  | YC-OC | 6.91 | 10.33 | 0.91 | 0.32 | 0.3850 | 3.59 | 10.47 | 0.49 | 0.19 | 0.6330 | 4.36 | 10.28 | 1.42 | 0.35 | 0.1860 |

| 8B #<br>Post-Learning |  | #Wake |  |  |  |  | #NREM |  |  |  |  | #REM |  |  |  |  |
| --- | --- | --- | --- | --- | --- | --- | --- | --- | --- | --- | --- | --- | --- | --- | --- | --- |
|  | Effect | Diff | DF | Tw | Ef.Size | p | Diff | DF | Tw | Ef.Size | p | Diff | DF | Tw | Ef.Size | p |
|  | YC-OC | 26.33 | 6.75 | 1.37 | 0.74 | 0.2150 | 46.81 | 9.23 | 2.84 | 0.74 | 0.0190* | 39.93 | 8.46 | 2.97 | 0.61 | 0.0170* |

| 8C Length<br>Post-Learning |  | Length Wake |  |  |  |  | Length NREM |  |  |  |  | Length REM |  |  |  |  |
| --- | --- | --- | --- | --- | --- | --- | --- | --- | --- | --- | --- | --- | --- | --- | --- | --- |
|  | Effect | Diff | DF | Tw | Ef.Size | p | Diff | DF | Tw | Ef.Size | p | Diff | DF | Tw | Ef.Size | p |
|  | YC-OC | 27.60 | 9.85 | 1.92 | 0.65 | 0.0850# | 43.11 | 10.97 | 2.09 | 0.61 | 0.0610# | 9.52 | 10.54 | 1.01 | 0.32 | 0.3370 |

Recovery
| 8D %<br>Recovery |  | %Wake |  |  |  | %NREM |  |  |  | %REM |  |  |  |
| --- | --- | --- | --- | --- | --- | --- | --- | --- | --- | --- | --- | --- | --- |
|  | Effect | DF | Fw | Ef.Size | p | DF | Fw | Ef.Size | p | DF | Fw | Ef.Size | p |
|  | Age | 1 | 0.33 | 0.08 | 0.5740 | 1 | 0.09 | 0.05 | 0.7650 | 1 | 0.84 | 0.23 | 0.3690 |
|  | Sleep | 1 | 3.02 | 0.43 | 0.099# | 1 | 4.65 | 0.36 | 0.0480* | 1 | 0.76 | 0.14 | 0.3920 |
|  | Age*Sleep | 1 | 0.04 | 0.08 | 0.8491 | 1 | 0.06 | 0.07 | 0.7369 | 1 | 0.08 | 0.16 | 0.6884 |

| 8E #<br>Recovery |  | #Wake |  |  |  | #NREM |  |  |  | #REM |  |  |  |
| --- | --- | --- | --- | --- | --- | --- | --- | --- | --- | --- | --- | --- | --- |
|  | Effect | DF | Fw | Ef.Size | p | DF | Fw | Ef.Size | p | DF | Fw | Ef.Size | p |
|  | Age | 1 | 1.23 | 0.20 | 0.2830 | 1 | 1.06 | 0.38 | 0.3210 | 1 | 0.40 | 0.15 | 0.5380 |
|  | Sleep | 1 | 1.99 | 0.28 | 0.1750 | 1 | 5.34 | 0.54 | 0.0350* | 1 | 0.02 | 0.08 | 0.8840 |
|  | Age*Sleep | 1 | 0.05 | 0.03 | 0.8191 | 1 | 0.32 | 0.29 | 0.0252* | 1 | 0.36 | 0.60 | 0.0468* |

| 8E #<br>Recovery<br>Age*Sleep Interaction |  | #NREM |  |  |  |  | #REM |  |  |  |  |
| --- | --- | --- | --- | --- | --- | --- | --- | --- | --- | --- | --- |
|  | Simple Effect | Diff | DF | Tw | Ef.Size | p | Diff | DF | Tw | Ef.Size | p |
|  | C-SR on Young | 23.71 | 8.88 | 1.04 | 0.30 | 0.3280 | 16.86 | 9.56 | 1.41 | 0.52 | 0.1910 |
|  | C-SR on Old | 38.33 | 5.66 | 2.73 | 0.79 | 0.0360* | 13.83 | 9.83 | 0.84 | 0.37 | 0.4220 |
|  | Y-O on Control | 6.50 | 10.10 | 0.27 | 0.18 | 0.7970 | 8.93 | 10.79 | 0.59 | 0.25 | 0.5650 |
|  | Y-O on SR | 21.12 | 7.33 | 1.94 | 0.58 | 0.0920# | 21.76 | 7.25 | 1.58 | 0.59 | 0.1570 |

| 8F Length Recovery |  | Length Wake |  |  |  | Length NREM |  |  |  | Length REM |  |  |  |
| --- | --- | --- | --- | --- | --- | --- | --- | --- | --- | --- | --- | --- | --- |
|  | Effect | DF | Fw | Ef.Size | p | DF | Fw | Ef.Size | p | DF | Fw | Ef.Size | p |
|  | Age | 1 | 0.40 | 0.15 | 0.5350 | 1 | 0.17 | 0.23 | 0.6900 | 1 | 1.36 | 0.28 | 0.2580 |
|  | Sleep | 1 | 0.02 | 0.05 | 0.8990 | 1 | 7.53 | 0.62 | 0.0130* | 1 | 4.19 | 0.43 | 0.0540# |
|  | Age*Sleep | 1 | 0.01 | 0.11 | 0.9674 | 1 | 0.06 | 0.08 | 0.7267 | 1 | 0.34 | 0.57 | 0.0591# |

Relative Power Analysis: Figure 6G-J
| 6G RDP Post-Learning | Effect | Diff | DF | Tw | Ef.Size | p |
| --- | --- | --- | --- | --- | --- | --- |
|  | YC-OC | 0.04 | 9.83 | 1.12 | 0.32 | 0.2906 |

| 6H RDP Recovery | Effect | DF | Fw | Ef.Size | p |
| --- | --- | --- | --- | --- | --- |
|  | Age | 1 | 0.79 | 0.14 | 0.3788 |
|  | Sleep | 1 | 5.17 | 0.52 | 0.0291* |
|  | Age*Sleep | 1 | 0.43 | 0.39 | 0.5177 |

| 6I RTP Post-Learning | Effect | Diff | DF | Tw | Ef.Size | p |
| --- | --- | --- | --- | --- | --- | --- |
|  | YC-OC | 0.02 | 10.98 | 0.91 | 0.27 | 0.3827 |

| 6J RTP Recovery | Effect | DF | Fw | Ef.Size | p |
| --- | --- | --- | --- | --- | --- |
|  | Age | 1 | 1.66 | 0.42 | 0.2061 |
|  | Sleep | 1 | 4.45 | 0.44 | 0.0421* |
|  | Age*Sleep | 1 | 2.73 | 0.68 | 0.1076 |

| Data not shown<br>RDP Normalized by REM | Effect | DF | Fw | Ef.Size | p |
| --- | --- | --- | --- | --- | --- |
|  | Age | 1 | 0.66 | 0.19 | 0.4300 |
|  | Sleep | 1 | 2.56 | 0.37 | 0.1290 |
|  | Age*Sleep | 1 | 0.35 | 0.16 | 0.5630 |

| Wake Post-Learning<br>Sleep restriction period<br>Data not shown | Effect | DF | Fw | Ef.Size | p |
| --- | --- | --- | --- | --- | --- |
|  | Age | 1 | 1.51 | 0.04 | 0.2838 |
|  | Band | 3 | 42.38 | 0.78 | 0.0150* |
|  | Age*Band | 3 | 1.15 | 0.43 | 0.5175 |

Spindle Analysis: Figure 7
| 9B Post-Learning<br>Average spindle count | Effect | Diff | DF | Tw | Ef.Size | p |
| --- | --- | --- | --- | --- | --- | --- |
|  | YC-OC | 4.33 | 10.99 | 0.52 | 0.14 | 0.6150 |

| 9B Post-Learning<br>Spindle count per Hour | Effect | DF | Fw | Ef.Size | p |
| --- | --- | --- | --- | --- | --- |
|  | Age | 1 | 0.86 | 0.02 | 0.3809 |
|  | Hour | 4 | 8.73 | 0.41 | 0.0080* |
|  | Age*Hour | 4 | 4.35 | 0.43 | 0.0459* |

| 9B Post-Learning<br>Spindle count per Hour<br>Age*Hour Interaction | Simple Effect | Diff | DF | Tw | Ef.Size | p |
| --- | --- | --- | --- | --- | --- | --- |
|  | YC- OC on Hr 1 | 128.83 | 10.06 | 4.18 | 0.82 | 0.0019* |
|  | YC- OC on Hr 2 | 15.45 | 8.23 | 0.34 | 0.10 | 0.7408 |
|  | YC- OC on Hr 3 | 35.71 | 6.96 | 1.01 | 0.33 | 0.3446 |
|  | YC- OC on Hr 4 | 40.43 | 10.24 | 1.29 | 0.38 | 0.2248 |
|  | YC- OC on Hr 5 | 14.36 | 7.36 | 0.36 | 0.11 | 0.7278 |

| 9C Post-Learning<br>Spindle duration | Effect | Diff | DF | Tw | Ef.Size | p |
| --- | --- | --- | --- | --- | --- | --- |
|  | YC-OC | 0.0001 | 9 | 0.004 | 0.05 | 0.9970 |

| 9C Post-Learning<br>Spindle duration per Hour | Effect | DF | Fw | Ef.Size | p |
| --- | --- | --- | --- | --- | --- |
|  | Age | 1 | 0.06 | 0.16 | 0.8193 |
|  | Hour | 4 | 0.64 | 0.20 | 0.6530 |
|  | Age*Hour | 4 | 1.15 | 0.45 | 0.4122 |

| 9D Post-Learning<br>Spindle frequency | Effect | Diff | DF | Tw | Ef.Size | p |
| --- | --- | --- | --- | --- | --- | --- |
|  | YC-OC | 0.0006 | 8.19 | 0.01 | 0.08 | 0.9957 |

| 9D Post-Learning<br>Spindle frequency per Hour | Effect | DF | Fw | Ef.Size | p |
| --- | --- | --- | --- | --- | --- |
|  | Age | 1 | 0.22 | 0.20 | 0.6511 |
|  | Hour | 4 | 0.81 | 0.19 | 0.6359 |
|  | Age*Hour | 4 | 0.34 | 0.13 | 0.8854 |

| 9E Recovery<br>Average spindle count | Effect | DF | Fw | Ef.Size | p |
| --- | --- | --- | --- | --- | --- |
|  | Age | 1 | 3.59 | 0.34 | 0.0750# |
|  | Sleep | 1 | 3.47 | 0.26 | 0.0800# |
|  | Age*Sleep | 1 | 1 | 0.61 | 0.0200* |

| **9E Recovery**<br>**Average spindle count**<br>Age*Sleep Interaction | Simple Effect | Diff | DF | Tw | Ef.Size | p |
| --- | --- | --- | --- | --- | --- | --- |
|  | C-SR on Young | 4.53 | 7.81 | 0.46 | 0.14 | 0.6560 |
|  | C-SR on Old | 29.10 | 8.85 | 3.29 | 0.58 | 0.0100* |
|  | Y-O on Control | 4.31 | 10.83 | 0.37 | 0.15 | 0.7190 |
|  | Y-O on SR | 29.32 | 9.37 | 4.77 | 0.76 | 0.0010* |

| **9E Recovery**<br>**Spindle count per Hour** | Effect | DF | Fw | p |
| --- | --- | --- | --- | --- |
|  | Age | 1 | 1.06 | 0.3077 |
|  | Sleep | 1 | 0.55 | 0.4626 |
|  | Hour | 4 | 0.90 | 0.0003* |
|  | Age*Sleep | 1 | 3.04 | 0.0000* |
|  | Age*Hour | 4 | 0.23 | 0.1078 |
|  | Sleep*Hour | 4 | 0.04 | 0.8312 |
|  | Age*Sleep*Hour | 4 | 0.07 | 0.6324 |

| **9E Recovery**<br>**Spindle count per Hour**<br>Age*Sleep Interaction | Simple Effect | Diff | DF | Tw | Ef.Size | p |
| --- | --- | --- | --- | --- | --- | --- |
|  | C-SR on Young | 33.24 | 37.64 | 0.43 | 0.14 | 0.6683 |
|  | C-SR on Old | 127.50 | 30.19 | 1.39 | 0.58 | 0.1752 |
|  | Y-O on Control | 20.31 | 34.13 | 0.21 | 0.10 | 0.8314 |
|  | Y-O on SR | 140.43 | 31.17 | 1.91 | 0.76 | 0.0651# |

| **9E Recovery**<br>**Spindle count per Hour**<br>Hour main effect | Main Effect | Diff | Zw | Ef.Size | p |
| --- | --- | --- | --- | --- | --- |
|  | Hr 1- Hr 2 | 169.83 | 2.31 | 0.19 | 0.0689# |
|  | Hr 2- Hr 3 | 98.33 | 0.93 | 0.34 | 0.3929 |
|  | Hr 3- Hr 4 | 109.67 | 0.90 | 0.32 | 0.4093 |
|  | Hr 4- Hr 5 | 97.67 | 3.05 | 0.17 | 0.0300* |

| **9F Recovery**<br>**Spindle duration** | Effect | DF | Fw | Ef.Size | p |
| --- | --- | --- | --- | --- | --- |
|  | Age | 1 | 0.23 | 0.08 | 0.6430 |
|  | Sleep | 1 | 0.30 | 0.09 | 0.5910 |
|  | Age*Sleep | 1 | 1.00 | 0.25 | 0.6580 |

| **9F Recovery**<br>**Spindle duration per Hour** | Effect | DF | Fw | p |
| --- | --- | --- | --- | --- |
|  | Age | 1 | 0.37 | 0.5420 |
|  | Sleep | 1 | 0.00 | 0.9740 |
|  | Hour | 4 | 0.37 | 0.8320 |
|  | Age*Sleep | 1 | 1.30 | 0.2550 |
|  | Age*Hour | 4 | 0.12 | 0.9770 |
|  | Sleep*Hour | 4 | 0.26 | 0.9010 |
|  | Age*Sleep*Hour | 4 | 0.08 | 0.9880 |
| **9G Recovery**<br>**Spindle frequency** | Effect | DF | Fw | Ef.Size | p |
|  | Age | 1 | 10.32 | 0.55 | 0.0040* |
|  | Sleep | 1 | 0.00 | 0.05 | 0.9980 |
|  | Age*Sleep | 1 | 0.37 | 0.54 | 0.0473* |

|  |  |  |  |  |  |  |
| --- | --- | --- | --- | --- | --- | --- |
| **9G Recovery**<br>**Spindle frequency**<br>Age*Sleep Interaction | Simple Effect | Diff | DF | Tw | Ef.Size | p |
|  | C-SR on Young | 0.09 | 11.92 | 1.27 | 0.36 | 0.2296 |
|  | C-SR on Old | 0.09 | 9.56 | 1.56 | 0.47 | 0.1519 |
|  | Y-O on Control | 0.06 | 11.00 | 0.87 | 0.31 | 0.4049 |
|  | Y-O on SR | 0.24 | 10.29 | 3.73 | 0.83 | 0.0037* |

|  |  |  |  |  |
| --- | --- | --- | --- | --- |
| **9G Recovery**<br>**Spindle frequency per Hour** | Effect | DF | Fw | p |
|  | Age | 1 | 7.94 | 0.0050* |
|  | Sleep | 1 | 0.00 | 0.9600 |
|  | Hour | 4 | 0.17 | 0.9600 |
|  | Age*Sleep | 1 | 2.91 | 0.0900# |
|  | Age*Hour | 4 | 0.02 | 0.9999 |
|  | Sleep*Hour | 4 | 0.24 | 0.9200 |
|  | Age*Sleep*Hour | 4 | 0.09 | 0.9999 |

## References

1. Abel, T., Havekes, R., Saletin, J.M., and Walker, M.P. (2013). Sleep, plasticity and memory from molecules to whole-brain networks. Curr Biol 23, R774–788.

2. Algina, J., and Olejnik, S.F. (1984). Implementing the Welch-James procedure with factorial design. Educational and Psychological Measurement 44, 39–48.

3. Altena, E., Ramautar, J.R., Van Der Werf, Y.D., and Van Someren, E.J. (2010). Do sleep complaints contribute to age-related cognitive decline? Prog Brain Res 185, 181–205.

4. Antony, J.W., Schonauer, M., Staresina, B.P., and Cairney, S.A. (2019). Sleep Spindles and Memory Reprocessing. Trends Neurosci 42, 1–3.

5. Bailey, C.H., Kandel, E.R., and Si, K. (2004). The persistence of long-term memory: a molecular approach to self-sustaining changes in learning-induced synaptic growth. Neuron 44, 49–57.

6. Barnes, C.A., Suster, M.S., Shen, J., and McNaughton, B.L. (1997). Multistability of cognitive maps in the hippocampus of old rats. Nature 388, 272–275.

7. Binder, S., Baier, P.C., Molle, M., Inostroza, M., Born, J., and Marshall, L. (2012). Sleep enhances memory consolidation in the hippocampus-dependent object-place recognition task in rats. Neurobiol Learn Mem 97, 213–219.

8. Buzsaki, G. (2015). Hippocampal sharp wave-ripple: A cognitive biomarker for episodic memory and planning. Hippocampus 25, 1073–1188.

9. Cam, E., Gao, B., Imbach, L., Hodor, A., and Bassetti, C.L. (2013). Sleep deprivation before stroke is neuroprotective: a pre-ischemic conditioning related to sleep rebound. Exp Neurol 247, 673–679.

10. Castello-Domenech, A.B., Del Valle, V.I., Fernandez-Garrido, J., Martinez-Martinez, M., and Cauli, O. (2016). Sleep Alterations in Non-demented Older Individuals: The Role of Cortisol. Endocr Metab Immune Disord Drug Targets 16, 174–180.

11. Clemens, Z., Fabo, D., and Halasz, P. (2006). Twenty-four hours retention of visuospatial memory correlates with the number of parietal sleep spindles. Neurosci Lett 403, 52–56.

12. Clemens, Z., Molle, M., Eross, L., Jakus, R., Rasonyi, G., Halasz, P., and Born, J. (2011). Fine-tuned coupling between human parahippocampal ripples and sleep spindles. Eur J Neurosci 33, 511–520.

13. Coenen, A.M., and van Luijtelaar, E.L. (1985). Stress induced by three procedures of deprivation of paradoxical sleep. Physiol Behav 35, 501–504.

14. Cohen, S.J., Munchow, A.H., Rios, L.M., Zhang, G., Asgeirsdottir, H.N., and Stackman, R.W., Jr. (2013). The rodent hippocampus is essential for nonspatial object memory. Curr Biol 23, 1685–1690.

15. Davis, C.J., Clinton, J.M., Jewett, K.A., Zielinski, M.R., and Krueger, J.M. (2011). Delta wave power: an independent sleep phenotype or epiphenomenon? J Clin Sleep Med 7, S16–18.

16. de Bruin, E.J., van Run, C., Staaks, J., and Meijer, A.M. (2017). Effects of sleep manipulation on cognitive functioning of adolescents: A systematic review. Sleep Med Rev 32, 45–57.

17. de Lavilleon, G., Lacroix, M.M., Rondi-Reig, L., and Benchenane, K. (2015). Explicit memory creation during sleep demonstrates a causal role of place cells in navigation. Nat Neurosci 18, 493–495.

18. Dispersyn, G., Sauvet, F., Gomez-Merino, D., Ciret, S., Drogou, C., Leger, D., Gallopin, T., and Chennaoui, M. (2017). The homeostatic and circadian sleep recovery responses after total sleep deprivation in mice. J Sleep Res 26, 531–538.

19. Dix, S.L., and Aggleton, J.P. (1999). Extending the spontaneous preference test of recognition: evidence of object-location and object-context recognition. Behav Brain Res 99, 191–200.

20. Drieu, C., Todorova, R., and Zugaro, M. (2018). Nested sequences of hippocampal assemblies during behavior support subsequent sleep replay. Science 362, 675–679.

21. Espiritu, J.R. (2008). Aging-related sleep changes. Clin Geriatr Med 24, 1–14

22. v. Fernandez, L.M.J., and Luthi, A. (2020). Sleep Spindles: Mechanisms and Functions. Physiol Rev 100, 805–868.

23. Gais, S., and Born, J. (2004). Declarative memory consolidation: mechanisms acting during human sleep. Learn Mem 11, 679–685.

24. Gais, S., Lucas, B., and Born, J. (2006). Sleep after learning aids memory recall. Learn Mem 13, 259–262.

25. Graves, L.A., Heller, E.A., Pack, A.I., and Abel, T. (2003). Sleep deprivation selectively impairs memory consolidation for contextual fear conditioning. Learn Mem 10, 168–176.

26. Grigg-Damberger, M.M. (2012). The AASM Scoring Manual four years later. J Clin Sleep Med 8, 323–332.

27. Halassa, M.M., Florian, C., Fellin, T., Munoz, J.R., Lee, S.Y., Abel, T., Haydon, P.G., and Frank, M.G. (2009). Astrocytic modulation of sleep homeostasis and cognitive consequences of sleep loss. Neuron 61, 213–219.

28. Harmony, T. (2013). The functional significance of delta oscillations in cognitive processing. Front Integr Neurosci 7, 83.

29. Hasan, S., Dauvilliers, Y., Mongrain, V., Franken, P., and Tafti, M. (2012). Age-related changes in sleep in inbred mice are genotype dependent. Neurobiol Aging 33, 195 e113–126.

30. Hasselmo, M.E. (2006). The role of acetylcholine in learning and memory. Curr Opin Neurobiol 16, 710–715.

31. Hasselmo, M.E., and Stern, C.E. (2014). Theta rhythm and the encoding and retrieval of space and time. Neuroimage 85 *Pt* *2*, 656–666.

32. Havekes, R., Bruinenberg, V.M., Tudor, J.C., Ferri, S.L., Baumann, A., Meerlo, P., and Abel, T. (2014). Transiently increasing cAMP levels selectively in hippocampal excitatory neurons during sleep deprivation prevents memory deficits caused by sleep loss. J Neurosci 34, 15715–15721.

33. Huang, Y.L., Liu, R.Y., Wang, Q.S., Van Someren, E.J., Xu, H., and Zhou, J.N. (2002). Age-associated difference in circadian sleep-wake and rest-activity rhythms. Physiol Behav 76, 597–603.

34. Hutchison, I.C., and Rathore, S. (2015). The role of REM sleep theta activity in emotional memory. Front Psychol 6, 1439.

35. Hwaun, E., and Colgin, L.L. (2019). CA3 place cells that represent a novel waking experience are preferentially reactivated during sharp wave-ripples in subsequent sleep. Hippocampus.

36. Isomura, Y., Sirota, A., Ozen, S., Montgomery, S., Mizuseki, K., Henze, D.A., and Buzsaki, G. (2006). Integration and segregation of activity in entorhinal-hippocampal subregions by neocortical slow oscillations. Neuron 52, 871–882.

37. Johansen, S. (1980). The Welch-James approximation to the distribution of the residual sum of squares in a weighted linear regression. Biometrika 67, 85–92.

38. Joo, H.R., and Frank, L.M. (2018). The hippocampal sharp wave-ripple in memory retrieval for immediate use and consolidation. Nat Rev Neurosci 19, 744–757.

39. Kaushal, N., Nair, D., Gozal, D., and Ramesh, V. (2012). Socially isolated mice exhibit a blunted homeostatic sleep response to acute sleep deprivation compared to socially paired mice. Brain Res 1454, 65–79.

40. Keinath, A.T., Wang, M.E., Wann, E.G., Yuan, R.K., Dudman, J.T., and Muzzio, I.A. (2014). Precise spatial coding is preserved along the longitudinal hippocampal axis. Hippocampus 24, 1533–1548.

41. Kentros, C.G., Agnihotri, N.T., Streater, S., Hawkins, R.D., and Kandel, E.R. (2004). Increased attention to spatial context increases both place field stability and spatial memory. Neuron 42, 283–295.

42. Kolbe, T., Palme, R., Tichy, A., and Rulicke, T. (2015). Lifetime Dependent Variation of Stress Hormone Metabolites in Feces of Two Laboratory Mouse Strains. PLoS One 10, e0136112.

43. Krause, A.J., Simon, E.B., Mander, B.A., Greer, S.M., Saletin, J.M., Goldstein-Piekarski, A.N., and Walker, M.P. (2017). The sleep-deprived human brain. Nat Rev Neurosci 18, 404–418.

44. Larkin, M.C., Lykken, C., Tye, L.D., Wickelgren, J.G., and Frank, L.M. (2014). Hippocampal output area CA1 broadcasts a generalized novelty signal during an object-place recognition task. Hippocampus 24, 773–783.

45. Lee, A.K., and Wilson, M.A. (2002). Memory of sequential experience in the hippocampus during slow wave sleep. Neuron 36, 1183–1194.

46. Lester, A.W., Moffat, S.D., Wiener, J.M., Barnes, C.A., and Wolbers, T. (2017). The Aging Navigational System. Neuron 95, 1019–1035.

47. Leutgeb, S., Leutgeb, J.K., Barnes, C.A., Moser, E.I., McNaughton, B.L., and Moser, M.B. (2005). Independent codes for spatial and episodic memory in hippocampal neuronal ensembles. Science 309, 619–623.

48. Lister, J.P., and Barnes, C.A. (2009). Neurobiological changes in the hippocampus during normative aging. Arch Neurol 66, 829–833.

49. Liu, X., Ramirez, S., Redondo, R.L., and Tonegawa, S. (2014). Identification and Manipulation of Memory Engram Cells. Cold Spring Harb Symp Quant Biol 79, 59–65.

50. Martinez-Vargas, M., Estrada Rojo, F., Tabla-Ramon, E., Navarro-Arguelles, H., Ortiz- Lailzon, N., Hernandez-Chavez, A., Solis, B., Martinez Tapia, R., Perez Arredondo, A., Morales- Gomez, J., et al. (2012). Sleep deprivation has a neuroprotective role in a traumatic brain injury of the rat. Neurosci Lett 529, 118–122.

51. McKenzie, S., and Buzsaki, G. (2016). Hippocampal Mechanisms for the Segmentation of Space by Goals and Boundaries. In Micro-, Meso- and Macro-Dynamics of the Brain, G. Buzsaki, and Y. Christen, eds. (Cham (CH)), pp. 1–21.

52. Mendelson, W.B., and Bergmann, B.M. (1999). EEG delta power during sleep in young and old rats. Neurobiol Aging 20, 669–673.

53. Miller, D.B., and O’Callaghan, J.P. (2005). Aging, stress and the hippocampus. Ageing Res Rev 4, 123–140.

54. Mizumori, S.J. (2006). Hippocampal place fields: a neural code for episodic memory? Hippocampus 16, 685–690.

55. Morawska, M.M., Buchele, F., Moreira, C.G., Imbach, L.L., Noain, D., and Baumann, C.R. (2016). Sleep Modulation Alleviates Axonal Damage and Cognitive Decline after Rodent Traumatic Brain Injury. J Neurosci 36, 3422–3429.

56. Morgan, E., Schumm, L.P., McClintock, M., Waite, L., and Lauderdale, D.S. (2017). Sleep Characteristics and Daytime Cortisol Levels in Older Adults. Sleep 40.

57. Mumby, D.G., Gaskin, S., Glenn, M.J., Schramek, T.E., and Lehmann, H. (2002). Hippocampal damage and exploratory preferences in rats: memory for objects, places, and contexts. Learn Mem 9, 49–57.

58. Muzzio, I.A., Kentros, C., and Kandel, E. (2009a). What is remembered? Role of attention on the encoding and retrieval of hippocampal representations. J Physiol 587, 2837–2854.

59. Muzzio, I.A., Levita, L., Kulkarni, J., Monaco, J., Kentros, C., Stead, M., Abbott, L.F., and Kandel, E.R. (2009b). Attention enhances the retrieval and stability of visuospatial and olfactory representations in the dorsal hippocampus. PLoS Biol 7, e1000140.

60. Naidoo, N., Ferber, M., Master, M., Zhu, Y., and Pack, A.I. (2008). Aging impairs the unfolded protein response to sleep deprivation and leads to proapoptotic signaling. J Neurosci 28, 6539–6548.

61. Nishida, M., and Walker, M.P. (2007). Daytime naps, motor memory consolidation and regionally specific sleep spindles. PLoS One 2, e341.

62. O’Keefe, J., and Dostrovsky, J. (1971). The hippocampus as a spatial map. Preliminary evidence from unit activity in the freely-moving rat. Brain Res 34, 171–175.

63. Ognjanovski, N., Broussard, C., Zochowski, M., and Aton, S.J. (2018). Hippocampal Network Oscillations Rescue Memory Consolidation Deficits Caused by Sleep Loss. Cereb Cortex 28, 3711–3723.

64. Ognjanovski, N., Maruyama, D., Lashner, N., Zochowski, M., and Aton, S.J. (2014). CA1 hippocampal network activity changes during sleep-dependent memory consolidation. Front Syst Neurosci 8, 61.

65. Ognjanovski, N., Schaeffer, S., Wu, J., Mofakham, S., Maruyama, D., Zochowski, M., and Aton, S.J. (2017). Parvalbumin-expressing interneurons coordinate hippocampal network dynamics required for memory consolidation. Nat Commun 8, 15039.

66. Oh, H.J., Song, M., Kim, Y.K., Bae, J.R., Cha, S.Y., Bae, J.Y., Kim, Y., You, M., Lee, Y., Shim, J., et al. (2018). Age-Related Decrease in Stress Responsiveness and Proactive Coping in Male Mice. Front Aging Neurosci 10, 128.

67. Ohayon, M.M., Carskadon, M.A., Guilleminault, C., and Vitiello, M.V. (2004). Meta-analysis of quantitative sleep parameters from childhood to old age in healthy individuals: developing normative sleep values across the human lifespan. Sleep 27, 1255–1273.

68. Oliveira, A.M., Hawk, J.D., Abel, T., and Havekes, R. (2010). Post-training reversible inactivation of the hippocampus enhances novel object recognition memory. Learn Mem 17, 155–160.

69. Pace-Schott, E.F., and Spencer, R.M. (2015). Sleep-dependent memory consolidation in healthy aging and mild cognitive impairment. Curr Top Behav Neurosci 25, 307–330.

70. Palchykova, S., Winsky-Sommerer, R., Meerlo, P., Durr, R., and Tobler, I. (2006). Sleep deprivation impairs object recognition in mice. Neurobiol Learn Mem 85, 263–271.

71. Panagiotou, M., Vyazovskiy, V.V., Meijer, J.H., and Deboer, T. (2017). Differences in electroencephalographic non-rapid-eye movement sleep slow-wave characteristics between young and old mice. Sci Rep 7, 43656.

72. Patel, K.M., and Hoel, D.G. (1973). Nonparametric test for interaction in factorial experiments. Journal of teh American Statistical Association 68, 615–620.

73. Peyrache, A., Battaglia, F.P., and Destexhe, A. (2011). Inhibition recruitment in prefrontal cortex during sleep spindles and gating of hippocampal inputs. Proc Natl Acad Sci U S A 108, 17207–17212.

74. Poe, G.R. (2017). Sleep Is for Forgetting. J Neurosci 37, 464–473.

75. Powers, M.M., and Clark, G. (1955). An evaluation of cresyl echt violet acetate as a Nissl stain. Stain Technol 30, 83–88.

76. Prince, T.M., Wimmer, M., Choi, J., Havekes, R., Aton, S., and Abel, T. (2014). Sleep deprivation during a specific 3-hour time window post-training impairs hippocampal synaptic plasticity and memory. Neurobiol Learn Mem 109, 122–130.

77. Ranck, J.B., Jr. (1973). Studies on single neurons in dorsal hippocampal formation and septum in unrestrained rats. I. Behavioral correlates and firing repertoires. Exp Neurol 41, 461–531.

78. Rasch, B., and Born, J. (2013). About sleep’s role in memory. Physiol Rev 93, 681–766.

79. Rosenzweig, E.S., and Barnes, C.A. (2003). Impact of aging on hippocampal function: plasticity, network dynamics, and cognition. Prog Neurobiol 69, 143–179.

80. Rosinvil, T., Lafortune, M., Sekerovic, Z., Bouchard, M., Dube, J., Latulipe-Loiselle, A., Martin, N., Lina, J.M., and Carrier, J. (2015). Age-related changes in sleep spindles characteristics during daytime recovery following a 25-hour sleep deprivation. Front Hum Neurosci 9, 323.

81. Rowland, D.C., Yanovich, Y., and Kentros, C.G. (2011). A stable hippocampal representation of a space requires its direct experience. Proc Natl Acad Sci U S A 108, 14654–14658.

82. Rytkonen, K.M., Zitting, J., and Porkka-Heiskanen, T. (2011). Automated sleep scoring in rats and mice using the naive Bayes classifier. J Neurosci Methods 202, 60–64.

83. Saletin, J.M., Goldstein-Piekarski, A.N., Greer, S.M., Stark, S., Stark, C.E., and Walker, M.P. (2016). Human Hippocampal Structure: A Novel Biomarker Predicting Mnemonic Vulnerability to, and Recovery from, Sleep Deprivation. J Neurosci 36, 2355–2363.

84. Sawangjit, A., Oyanedel, C.N., Niethard, N., Salazar, C., Born, J., and Inostroza, M. (2018). The hippocampus is crucial for forming non-hippocampal long-term memory during sleep. Nature 564, 109–113.

85. Schimanski, L.A., and Barnes, C.A. (2010). Neural Protein Synthesis during Aging: Effects on Plasticity and Memory. Front Aging Neurosci 2.

86. Siapas, A.G., and Wilson, M.A. (1998). Coordinated interactions between hippocampal ripples and cortical spindles during slow-wave sleep. Neuron 21, 1123–1128.

87. Sirota, A., Csicsvari, J., Buhl, D., and Buzsaki, G. (2003). Communication between neocortex and hippocampus during sleep in rodents. Proc Natl Acad Sci U S A 100, 2065–2069.

88. Skaggs, W., McNaughton, B., Gothard, K., and Markus, E. (1993). An information-theoretic approach to deciphering the hippocampal code. In Advances in Neural Information Processing, S. Hanson, J. Cowan, and C. Giles, eds. (San Mateo, CA: Morgan Kaufmann), pp. 1030–1037.

89. Smith, C. (2001). Sleep states and memory processes in humans: procedural versus declarative memory systems. Sleep Med Rev 5, 491–506.

90. Smith, C., and Rose, G.M. (1996). Evidence for a paradoxical sleep window for place learning in the Morris water maze. Physiol Behav 59, 93–97.

91. Smith, D.M., and Mizumori, S.J. (2006). Hippocampal place cells, context, and episodic memory. Hippocampus 16, 716–729.

92. Sportiche, N., Suntsova, N., Methippara, M., Bashir, T., Mitrani, B., Szymusiak, R., and McGinty, D. (2010). Sustained sleep fragmentation results in delayed changes in hippocampal-dependent cognitive function associated with reduced dentate gyrus neurogenesis. Neuroscience 170, 247–258.

93. Squire, L.R., and Zola, S.M. (1998). Episodic memory, semantic memory, and amnesia. Hippocampus 8, 205–211.

94. Stickgold, R., and Walker, M.P. (2013). Sleep-dependent memory triage: evolving generalization through selective processing. Nat Neurosci 16, 139–145.

95. Taillard, J., Sagaspe, P., Berthomier, C., Brandewinder, M., Amieva, H., Dartigues, J.F., Rainfray, M., Harston, S., Micoulaud-Franchi, J.A., and Philip, P. (2019). Non-REM Sleep Characteristics Predict Early Cognitive Impairment in an Aging Population. Front Neurol 10, 197.

96. Tanaka, K.Z., He, H., Tomar, A., Niisato, K., Huang, A.J.Y., and McHugh, T.J. (2018). The hippocampal engram maps experience but not place. Science 361, 392–397.

97. Tartar, J.L., Ward, C.P., McKenna, J.T., Thakkar, M., Arrigoni, E., McCarley, R.W., Brown, R.E., and Strecker, R.E. (2006). Hippocampal synaptic plasticity and spatial learning are impaired in a rat model of sleep fragmentation. Eur J Neurosci 23, 2739–2748.

98. Team, R.C. (2020). R: A language and environment for statistical computing, R.F.f.S. Computing, ed. (Vienna, Austria).

99. Tobler, I., Murison, R., Ursin, R., Ursin, H., and Borbely, A.A. (1983). The effect of sleep deprivation and recovery sleep on plasma corticosterone in the rat. Neurosci Lett 35, 297–300.

100. Tononi, G., and Cirelli, C. (2006). Sleep function and synaptic homeostasis. Sleep Med Rev 10, 49–62.

101. Tononi, G., and Cirelli, C. (2014). Sleep and the price of plasticity: from synaptic and cellular homeostasis to memory consolidation and integration. Neuron 81, 12–34.

102. Ulrich, D. (2016). Sleep Spindles as Facilitators of Memory Formation and Learning. Neural Plast 2016, 1796715.

103. Uygun, D.S., Katsuki, F., Bolortuya, Y., Aguilar, D.D., McKenna, J.T., Thankachan, S., McCarley, R.W., Basheer, R., Brown, R.E., Strecker, R.E., et al. (2019). Validation of an automated sleep spindle detection method for mouse electroencephalography. Sleep 42.

104. Wang, M.E., Wann, E.G., Yuan, R.K., Ramos Alvarez, M.M., Stead, S.M., and Muzzio, I.A. (2012). Long-term stabilization of place cell remapping produced by a fearful experience. J Neurosci 32, 15802–15814.

105. Wang, M.E., Yuan, R.K., Keinath, A.T., Ramos Alvarez, M.M., and Muzzio, I.A. (2015). Extinction of Learned Fear Induces Hippocampal Place Cell Remapping. J Neurosci 35, 9122–9136.

106. Ward, C.P., McCarley, R.W., and Strecker, R.E. (2009a). Experimental sleep fragmentation impairs spatial reference but not working memory in Fischer/Brown Norway rats. J Sleep Res 18, 238–244.

107. Ward, C.P., McCoy, J.G., McKenna, J.T., Connolly, N.P., McCarley, R.W., and Strecker, R.E. (2009b). Spatial learning and memory deficits following exposure to 24 h of sleep fragmentation or intermittent hypoxia in a rat model of obstructive sleep apnea. Brain Res 1294, 128–137.

108. Wennberg, A.M., Canham, S.L., Smith, M.T., and Spira, A.P. (2013). Optimizing sleep in older adults: treating insomnia. Maturitas 76, 247–252.

109. Wilcox, R.R. (2012). Introduction to robust estimation hypothesis testing (San Diego, CA: Academic Press).

110. Wilson, M., Permito, R., English, A., Albritton, S., Coogle, C., and Van Dongen, H.P.A. (2019). Performance and sleepiness in nurses working 12-h day shifts or night shifts in a community hospital. Accid Anal Prev 126, 43–46.

111. Wimmer, M.E., Hernandez, P.J., Blackwell, J., and Abel, T. (2012). Aging impairs hippocampus-dependent long-term memory for object location in mice. Neurobiol Aging 33, 2220–2224.

112. Zheng, C., Bieri, K.W., Hwaun, E., and Colgin, L.L. (2016). Fast Gamma Rhythms in the Hippocampus Promote Encoding of Novel Object-Place Pairings. eNeuro 3.

